# Single-cell transcriptomics reveals striking heterogeneity and functional organization of dendritic and monocytic cells in the bovine mesenteric lymph node

**DOI:** 10.1101/2022.10.24.513393

**Authors:** G.T. Barut, M.C. Kreuzer, R. Bruggmann, A. Summerfield, S.C. Talker

## Abstract

Dendritic and monocytic cells co-operate to initiate and shape adaptive immune responses in secondary lymphoid tissue. The complexity of this system is poorly understood, also because of the high phenotypic and functional plasticity of monocytic cells. We have sequenced mononuclear phagocytes in mesenteric lymph nodes (LN) of three adult cows at the single-cell level, revealing ten dendritic-cell (DC) clusters and seven monocyte/macrophage clusters with clearly distinct transcriptomic profiles. Among DC, we defined LN-resident subsets and their progenitors, as well as subsets of highly activated migratory DC differing in transcript levels for T-cell attracting chemokines. Our analyses also revealed a potential differentiation path for cDC2, resulting in a cluster of inflammatory cDC2 with close transcriptional similarity to putative DC3 and monocyte-derived DC. Monocytes and macrophages displayed sub-clustering mainly driven by pro- or anti-inflammatory expression signatures, including a small cluster of cycling, presumably self-renewing, macrophages.

With this transcriptomic snapshot of LN-derived mononuclear phagocytes, we reveal functional properties and differentiation trajectories in a “command center of immunity” that are likely to be conserved across species.

## INTRODUCTION

Dendritic cells (DC) are known as central instructors of adaptive immunity (1), which is initiated in specialized areas of secondary lymphoid tissues (2). It is widely accepted that also monocytic cells contribute to shape adaptive immune responses (3, 4). Being about ten times more frequent than DC in peripheral blood, monocytes fulfill multiple functions in the induction and resolution of inflammation (5), and have also been implicated in antigen presentation in lymph nodes (6).

Bona fide DCs can be delineated from other mononuclear phagocytes by their surface expression of Flt3 receptor tyrosine kinase, a hematopoietic cytokine receptor essential for DC differentiation (7–11). Distinction of three DC subsets (cDC1, cDC2, pDC) is well established and based on differences in ontogeny, expression of key genes, phenotype, and function (12–14). Murine studies have suggested that cDC1 and cDC2 are specialized to induce CD8-T cell/Th1 responses and Th2/Th17 responses, respectively (15, 16). As such, murine cDC1 were described to be especially efficient at cross-presentation (17). Recent findings have strengthened the idea that also pDC engage in T-cell stimulation, notably following initial IFN-I production (18). To identify DC subsets in non-mouse / non-human species (namely pig, horse, and cow), we have previously analyzed expression of key genes that were found to be evolutionarily conserved in DC subsets, including essential transcription factors and *FLT3* (9–11).

Across species, monocyte subsets are less well defined and most likely represent a differentiation continuum rather than developmentally distinct populations (19). Nevertheless, in humans and cattle, nonclassical and intermediate monocytes (ncM and intM) can be distinguished from classical monocytes (cM) based on expression of CD14/CD16 and certain key genes such as *CCR2* (cM) and *NR4A1*/*CX3CR1* (ncM and intM) (11, 20–22). Notably, we recently demonstrated that bovine monocyte subsets appear to be very similar to their human counterparts (22, 23), with transcriptomic analyses suggesting a role of ncM in anti-viral immunity and a specialization of intM in antigen presentation. In tissues, however, the subset identity of monocytes gets blurred, as they are differentiating towards monocyte-derived DC (moDC) and monocyte-derived macrophages, the latter of which can be transcriptionally very similar to long-lived and self-renewing tissue-resident macrophages of embryonic origin (3, 24, 25).

Single-cell RNA sequencing (scRNA-seq) has enabled a relatively unbiased view on immune-cell composition in complex tissues (26), revealing for example considerable heterogeneity within human cDC2, including an inflammatory DC subset with close similarity to moDC (27–30), and contributing to the recent discovery of the separate DC lineage DC3 – cells described as phenotypic and functional intermediates between cDC2 and monocytes (31, 32).

Moreover, scRNA-seq enables high-level comparative immunology through in-depth analysis of non-model species, promising exciting insights into basic and evolutionarily conserved functions of immune cells. Having previously performed detailed transcriptomic analyses of dendritic and monocytic cells isolated from blood of cattle, both *ex vivo* and following *in-vitro* stimulation (11, 22, 33), we provide here the first scRNA-seq analysis of the mononuclear phagocyte compartment in bovine secondary lymphoid tissue. With the mesenteric lymph node we have chosen a secondary lymphoid tissue that is expected to contain a considerable fraction of activated immune cells under steady-state conditions – giving us the opportunity to study the mononuclear phagocyte system at its best – where antigens are presented, T cells get instructed, and apoptotic cells are disposed of.

## MATERIALS AND METHODS

### Isolation of bovine mesenteric-lymph-node cells

Mesenteric lymph nodes (mesLN) draining the small intestine were collected from three cows that were slaughtered at a nearby butchery in three consecutive weeks (MLN2306: RedHolstein, 3.5 years; MLN3006: Simmental, 8.5 years; MLN0707: Holstein, 4 years), and immediately placed into ice-cold PBS containing 1 mM EDTA (Invitrogen, ThermoFisher, Basel, Switzerland) (PBS-EDTA). After removal of fat and connective tissue, lymph nodes were cut into small pieces and minced using a gentleMACS Dissociator (Miltenyi Biotec Swiss AG, Solothurn, Switzerland). After a washing step with cold PBS-EDTA (300 x g, 10 min, 4°C), the cells were incubated in PBS containing 0.1 mg/mL DNase I (Worthington-Biochem, BioConcept, Basel, Switzerland), for 15 min at room temperature. Thereafter, cells were washed (300 x g, 10 min, 4°C), with DMEM supplemented with 5% FBS (Gibco, Life Technologies, Basel, Switzerland), and the suspension was passed through a cell strainer (70 μm). This washing step was repeated, and cells were resuspended in PBS-EDTA (room temperature) and layered onto Ficoll Paque (1.077 g/mL; GE Healthcare Europe GmbH). After centrifugation (800 x g, 25 min, 20°C), cells at the interface were collected and washed twice with cold PBS-EDTA (400 x g, 10 min, 4°C). Subsequently, the cells were counted and stained for fluorescence-activated cell sorting (FACS) as described below.

### Fluorescence-activated cell sorting (FACS) of Flt3^+^ and CD172a^high^ cells

Staining of isolated lymph-node cells was performed with 2 × 10^8^ cells in 50 mL Falcon tubes and encompassed three incubation steps, each for 20 min at 4°C (incubation volume 2mL). In-between incubation steps, cells were washed (300 x g, 10 min, 4°C) with 40 mL of BD Cell Wash (BD Biosciences, Allschwil, Switzerland). In order to block Fc receptors, cells were incubated with purified 50 μg/mL bovine IgG (Bethyl laboratories, Montgomery, USA), before adding CD172a (CC149, mIgG2b; Bio Rad, Cressier, Switzerland) and His-tagged Flt3L in a subsequent incubation step. Bovine His-tagged Flt3L (NCBI NM_181030.2) was produced as previously described (8, 10).

In the third and final step, cells were incubated with anti-IgG2b-AF647 (Molecular Probes, Thermo Fischer, Basel, Switzerland), anti-His-PE (mIgG1) (Miltenyi Biotec Swiss AG, Solothurn, Switzerland), and LIVE/DEAD™ Fixable Near-IR stain (Thermo Fisher Scientific, Basel, Switzerland). After a final washing step, cells were resuspended in BD Cell Wash and delivered to the Flow Cytometry and Cell Sorting Facility (FCCS) of the University of Bern, where 1×10^5^ viable cells (Flt3^+^ and/or CD172a^high^) were sorted using a MoFlo Astrios EQ cell sorter equipped with five lasers (Beckman Coulter Eurocenter SA, Nyon, Switzerland). After sorting, cells were spun down (300 x g, 10 min, 4 °C), re-suspended in 200 μl of PBS containing 0.4% BSA and viability was assessed microscopically using Trypan Blue staining. All three samples were confirmed to be free of visible debris and doublets and to have a viability > 95 %. Immediately after counting, cells were submitted to the Next Generation Sequencing Platform of the University of Bern for generation of 10x Genomics sequencing libraries and subsequent sequencing.

### Single-cell RNA sequencing (10x Genomics)

Library preparation was done in three consecutive weeks at the Next Generation Sequencing (NGS) Platform at the University of Bern. The three libraries were stored at minus 80 °C, before being pooled and sequenced in one run. GEM generation & barcoding, reverse transcription, cDNA amplification and 3’ gene expression library generation steps were all performed according to the Chromium Next GEM Single Cell 3’ Reagent Kits v3.1 User Guide (10x Genomics CG000204 Rev D) with all stipulated 10x Genomics reagents. Specifically, 33.0 μL of each cell suspension (500 cells/μL) and 10.2 μL of nuclease-free water were used for a targeted cell recovery of 10’000 cells. GEM generation was followed by a GEM-reverse transcription incubation, a clean-up step and 11 cycles of cDNA amplification. The resulting cDNA was evaluated for quantity and quality using a Thermo Fisher Scientific Qubit 4.0 fluorometer with the Qubit dsDNA HS Assay Kit (Thermo Fisher Scientific, Q32854) and an Advanced Analytical Fragment Analyzer System using a Fragment Analyzer NGS Fragment Kit (Agilent, DNF-473), respectively. Thereafter, 3′ gene expression libraries were constructed using a sample index PCR step of 13-15 cycles. At the end of the protocol, an additional 0.8x bead-based cleanup of the libraries was performed. The generated cDNA libraries were tested for quantity and quality using fluorometry and capillary electrophoresis as described above. The cDNA libraries were pooled and sequenced with a loading concentration of 300 pM, paired-end and single-indexed, on an illumina NovaSeq 6000 sequencer with a shared NovaSeq 6000 S2 Reagent Kit (100 cycles; illumina 20012862). The read set-up was as follows: read 1: 28 cycles, i7 index: 8 cycles, i5: 0 cycles and read 2: 91 cycles. The quality of the sequencing runs was assessed using illumina Sequencing Analysis Viewer (illumina version 2.4.7) and all base-call files were demultiplexed and converted into FASTQ files using illumina bcl2fastq conversion software v2.20. An average of 787,553,242 reads/library were obtained.

### Analysis of scRNA-seq data

Mapping and counting of the UMIs were performed using Cell Ranger (version 3.0.2, 10x Genomics) with the reference genome ARS-UCD1.2 from Ensembl to build the necessary index files. Subsequent analysis was performed in R (version 4.0.2) (34). The Scater package (version 1.14) (35) was used to assess the proportion of ribosomal and mitochondrial genes as well as the number of detected genes. Cells were considered as outliers and filtered out if the value of the proportion of expressed mitochondrial genes or the number detected genes deviated more than three median absolute deviations from the median across all cells. After quality control, the sample from MLN2306 retained 6604 cells, the sample from MLN3006 retained 3956 cells, and the sample from MLN0707 retained 3649 cells. Normalization between samples was done with the deconvolution method of Lun et al. (36) using the package Scran (version 1.14) (37). Samples were integrated with the FindIntegrationAnchors function of the package Seurat (version 3.1) based on the first 20 principal components (PCs) (38). Graph-based clustering was done with the FindNeighbors and FindClusters functions of the Seurat package using the first 40 PCs from the dimensionality reduction step. The Clustree package (version 0.4) (39) was used to determine the resolution resulting in clustering concurring with the presumed cell types, which was 1.2. In order to identify up- or down-regulated genes between clusters, FindAllMarkers was applied to the dataset. This function returns all differentially expressed genes per cluster. Clusters were then manually annotated on the basis of these marker genes. Upon publication of the final version, datasets will be available in the European Nucleotide Archive (ENA).

### Preparation of figures

Figures were prepared using FlowJo version 10 (FlowJo LLC, Ashland, OR), R version 4.1.1, and Inkscape (http://www.inkscape.org).

## RESULTS

### Distinct clustering of dendritic and monocytic cells

Single-cell transcriptomes were obtained from sorted Flt3^+^ and/or CD172a^high^ mesenteric-lymph-node cells of three healthy cows **(Figure 1A and Supplementary File 1)** and bioinformatically processed as outlined in Materials and Methods. The integrated dataset was used for further analyses. A resolution of 1.2 was chosen, resulting in 24 different clusters **(Figure 1B and Supplementary File 1)**. When visualizing the expression of the DC marker *FLT3* and the monocytic marker *CSF1R* in UMAP plots **(Figure 1C)**, we found that most of these 24 clusters could be grouped into major clusters of putative DC (c1, c3, c5, c6, c8, c9, c11, c12, c13, c15, c17, c20, c23) and putative monocytic cells (clusters c0, c2, c4, c7, c10, c14, c19, c21). Notably, clusters 3 and 15, both assigned to DC, contained a subset of cells expressing *CSF1R* and lacking *FLT3* expression.

**Figure 1.**
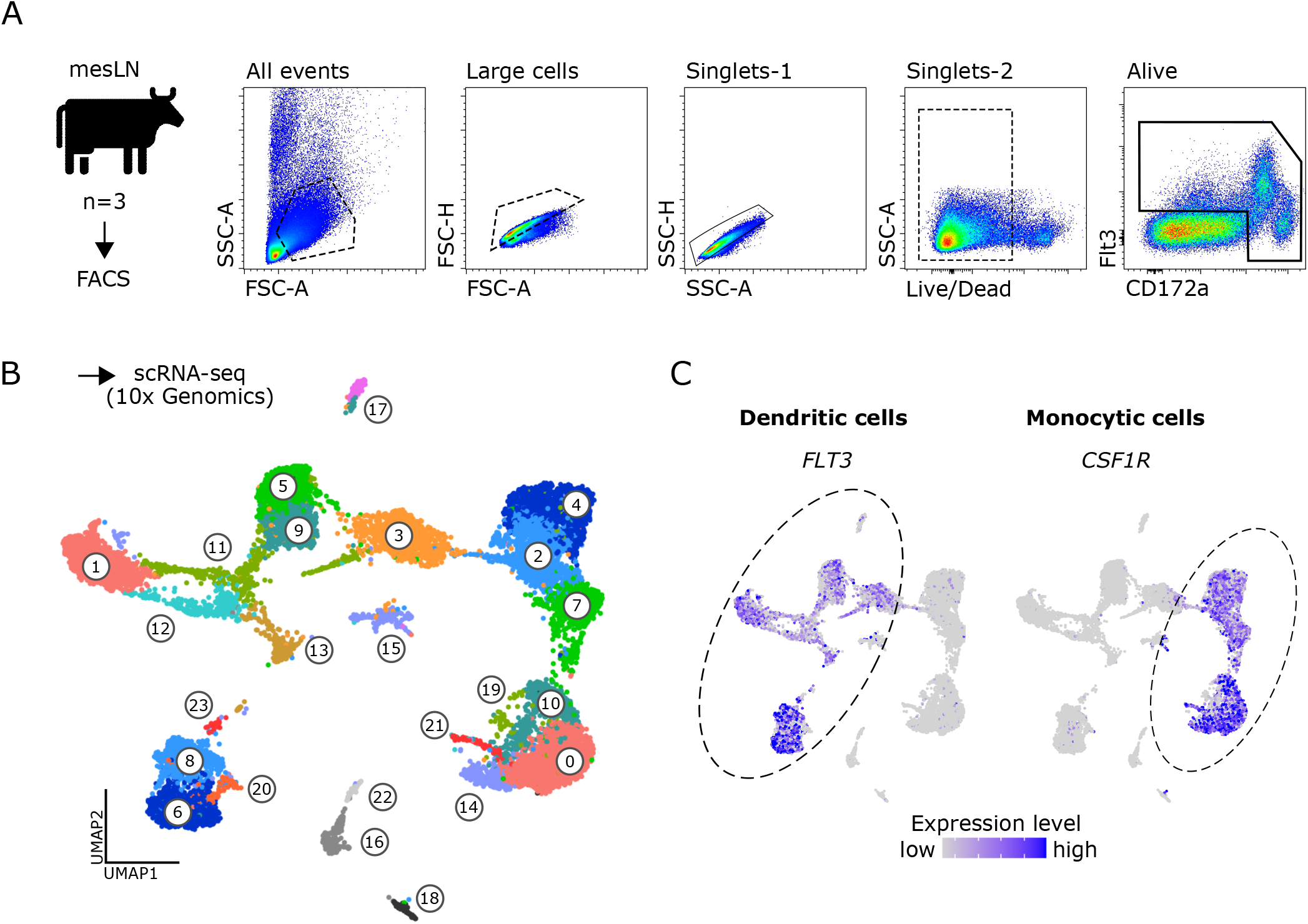
Single-cell RNA-seq of bovine mononuclear phagocytes. (A) Mononuclear phagocytes were sorted by FACS as CD172a^high^ and/or Flt3^+^ cells from mesenteric lymph nodes of three cows and subjected to 10x Genomics scRNA-seq. (B) Data of three animals were integrated and analyzed at a resolution of 1.2, resulting in 24 different clusters. (C) Visualization of *FLT3* and *CSF1R* in feature plots, revealing clusters of dendritic and monocytic cells, respectively.

Clusters lacking both *FLT3* and *CSF1R* expression, were found to express *CD79B* (c16 and c18) or *CD3E* (c22), alongside other B- and T-cell markers, respectively. Notably, c18 likely contained plasma cells, as indicated by the expression of *IRF4, PRDM1* (Blimp-1), *JCHAIN* and immunoglobulin genes. Notably these plasma cells expressed *CD27* and *TNFRSF17*, two surface molecules that could be targeted for flow cytometric detection of plasma cells in cattle.

Cluster-defining marker genes, as determined by seurat’s FindAllMarkers function, are listed in **Supplementary File 2**.

### Subset-specific gene transcription defines resident cDC1, cDC2 and pDC

In accordance with the phenotype and transcriptome of bona fide DC subsets recently identified in blood of cattle (11), distinct clusters of lymph-node-derived DC could be defined by expression of *ANPEP* (CD13), *FCER1A* and *CD4* **(Figure 2A)**.

**Figure 2.**
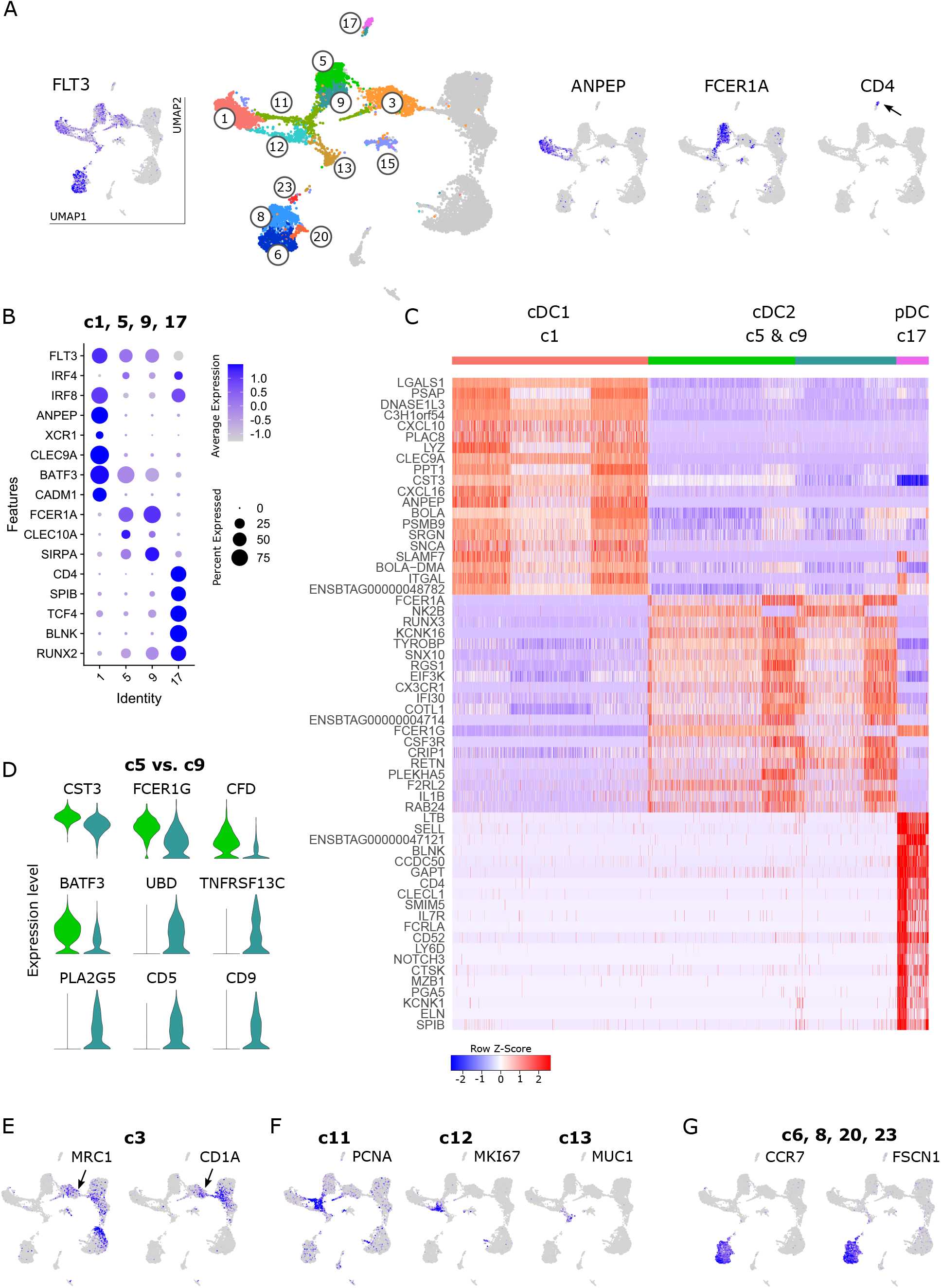
Dendritic cells. (A) Visualization of key genes in feature plots to identify clusters containing total DCs (*FLT3*), cDC1 (*ANPEP*, cluster 1), cDC2 (*FCER1A*, clusters 5 and 9) and pDC (*CD4*, cluster 17). (B) Dot plot of key subset-defining genes in selected DC clusters. (C) Heatmap of the top 20 (p_adj) differentially expressed genes in clusters identified as cDC1 (c1), cDC2 (c5 & c9) and pDC (c17). (D) Top genes differentially expressed between cDC2 clusters c5 and c9, visualized as violin plots (linear y-axis). (E-G) Expression of selected signature genes for DC clusters that could not be clearly assigned to a DC subset based on key gene expression.

As shown in **Figure 2B**, the co-expression of subset-specific key genes clearly confirmed cluster 1 as cDC1 (*ANPEP, XCR1, CLEC9A, BATF3, CADM1*), clusters 5 and 9 as cDC2 (*FCER1A, CLEC10A, SIRPA*), and cluster 17 as pDC (*CD4, SPIB, TCF4, BLNK, RUNX2*). A heatmap of the top 20 (adjusted p-value) differentially expressed genes between these clusters (c1, c5 & c9, c17) is shown in Figure 2C. The complete gene list is given in **Supplementary File 3**. Apart from the genes mentioned above, the top subset-specific genes included *LGALS1, PSAP, DNASE1L3, CXCL10, LYZ* for cDC1, *NK2B, RUNX3, CX3CR1* for cDC2, and *LTB, SELL, CLECL1* and *IL7R* for pDC.

The two cDC2 clusters (c5 and c9) differed in the expression of various immune-relevant genes **(Figure 2D and Supplementary File 4)**, such as *CST3, FCER1G, CFD*, and *BATF3* (higher expression in c5), and *UBD, TNFRSF13C, PLA2G5, CD5* and *CD9* (clearly expressed in c9, but almost absent from c5). Notably, these differences in CD5 and CD9 expression might be useful to distinguish bovine cDC2 subsets with flow cytometry.

Selected subset-specific key genes shown in **Figure 2B** were are also visualized for the complete dataset **(Supplementary File 5)**. Clusters 3, 6, 8, 13, 15, 20 and 23 lacked transcripts for most subset-defining marker genes listed above. Among DC, c3 exclusively contained cells expressing *MRC1* and *CD1A* **(Figure 2E)**. Clusters 11 and 12 appeared to contain both cDC1 and cDC2 and stood out by their high expression of cell cycle genes such as *PCNA* (c11), and *MKI67* (c12) **(Figure 2F)**. In fact, c11 and c12 predominantly expressed genes associated with the G1-S phase and the G2-M phase of the cell cycle, respectively **(Supplementary File 2)**. Furthermore, as presented in the UMAP projection **(Figure 2A)**, c13 conveyed the impression of giving rise to c11 and c12 that later diverge to meet cDC1 (c1) and cDC2 (c9). In cluster 13, however, we did not detect transcripts related to an active cell cycle. Instead, c13 stood out by high expression of *AATF* (cell-cycle control) as well as of *CD9, CD164L2, ANXA1, ANXA2*, and *CLEC10A*, and exclusive expression of *MUC1, RORC* and *SLC14A1* among other genes **(Figure 2F and Supplementary File 2)**.

Finally, despite their lack of DC-subset defining transcripts **(Supplementary File 5)**, clusters 6 and 8 showed by far the highest levels of *FLT3* expression, alongside exclusive expression of *CCR7* and *FSCN1* **(Figure 2G)**. In fact, their high levels of *CCR7* and *FSCN1* expression suggest that these cells comprise migratory DC that have recently migrated from intestinal tissue to the mesenteric lymph node.

### Subsets of CCR7^high^ migratory DC defined by chemokine expression

Cells in clusters 6, 8, 20 and 23, showing an isolated projection in the UMAP plot, stood out by expressing high levels of various activation- and maturation-related genes **(Figure 3A and Supplementary File 2)**. Over 1500 genes were exclusively expressed or significantly upregulated (p_adj < 0.05) in these migratory DC (c6 & c8 & c20 & c23) when compared against resident DC (c1 & c5 & c9 & c17) **(Supplementary File 6)**. This includes genes involved in cytoskeleton regulation (*FSCN1, SAMSN1, MARCKSL1*), T-cell or B-cell co-stimulation (*CD83, TNFSF13B, CD40*) and T-cell regulation (*IDO1, CD274, IL4I1*).

**Figure 3.**
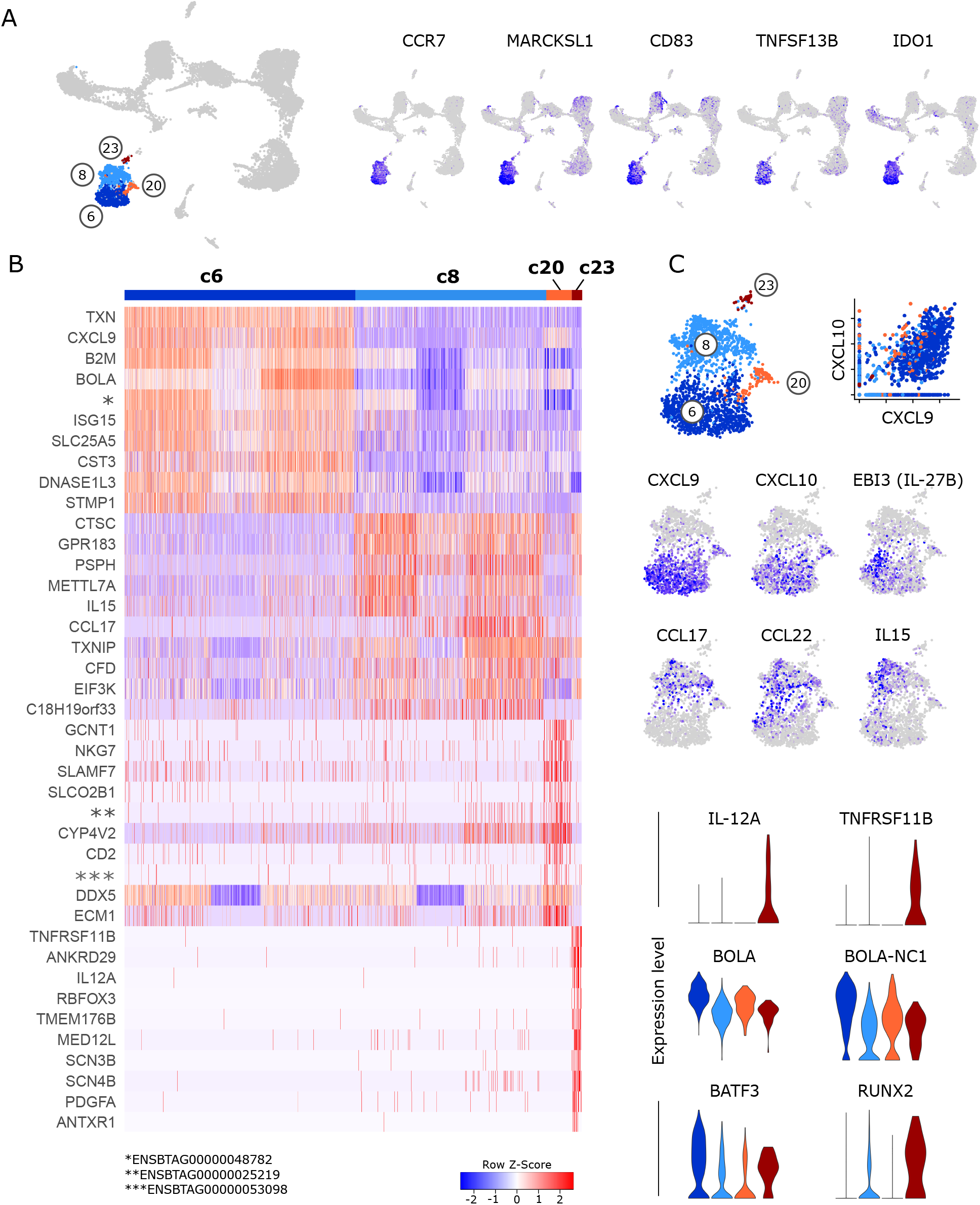
Migratory dendritic cells. (A) Feature plots illustrating high expression of selected activation-related genes in *CCR7*-expressing migratory DC (c6, c8, c20, c23). (B) Heatmap of the top 10 (p_adj) differentially expressed genes in clusters of *CCR7*-expressing DC. (C) Visualization of selected genes specifically expressed in clusters of *CCR7*-expressing DC.

Subclusters within *CCR7*^high^ migratory DC differed in the expression of T-cell attracting chemokines and T-cell activating cytokines **(Figure 3B and C)**, suggesting a division of labor between subsets of migratory DC (for complete gene lists see **Supplementary File 7**).

Cluster 6 stood out by high expression of T-cell attracting *CXCL9* and *CXCL10* (40, 41), alongside *ISG15* and MHC-I-associated genes. Notably, also *TXN* was specifically expressed in c6, encoding for thioredoxin, which has been suggested to regulate the Th1/Th2 balance, among other immuno-relevant effects (42). Moreover, *IL27* and *EBI3* (IL-27B) were specifically expressed in a subcluster of c6. Cells in cluster 8 were clearly enriched in transcripts for *CCL17* and *CCL22*, presumably attracting Th2 and Th17 cells as well as Tregs (43), and in *IL15* transcripts, promoting survival and proliferation of T cells and NK cells. Notably, the very small but distinct cluster 23, exclusively displayed expression of *IL12A* and *TNFRSF11B*. Although transcripts for most DC-lineage defining key genes were only weakly detected in these migratory DC clusters, overall gene expression would suggest that cDC2 are enriched in cluster 8 and that cDC1 are enriched in c6. Also the relatively high level of *BATF3, BOLA* and *BOLA-NC1* expression in cluster 6 would point towards cDC1, whereas the high level of *RUNX2* in cluster 23 may indicate the presence of pDC **(Figure 3C and Supplementary File 7)**.

### Co-clustering of inflammatory cDC2, monocyte-derived DC and putative DC3

Positioned in-between cDC2 (c5, c9) and monocytes (c2) in the UMAP projection, *FLT3*^+^ dendritic cells in cluster 3 expressed genes exclusively shared with either cDC2 (e.g. *F2RL2*) or monocytic cells (e.g. *MRC1*). Notably, a considerable fraction of cells in cluster 3 appeared to co-express these genes, including *FLT3* and *CSF1R* **(Figure 4A)**.

**Figure 4.**
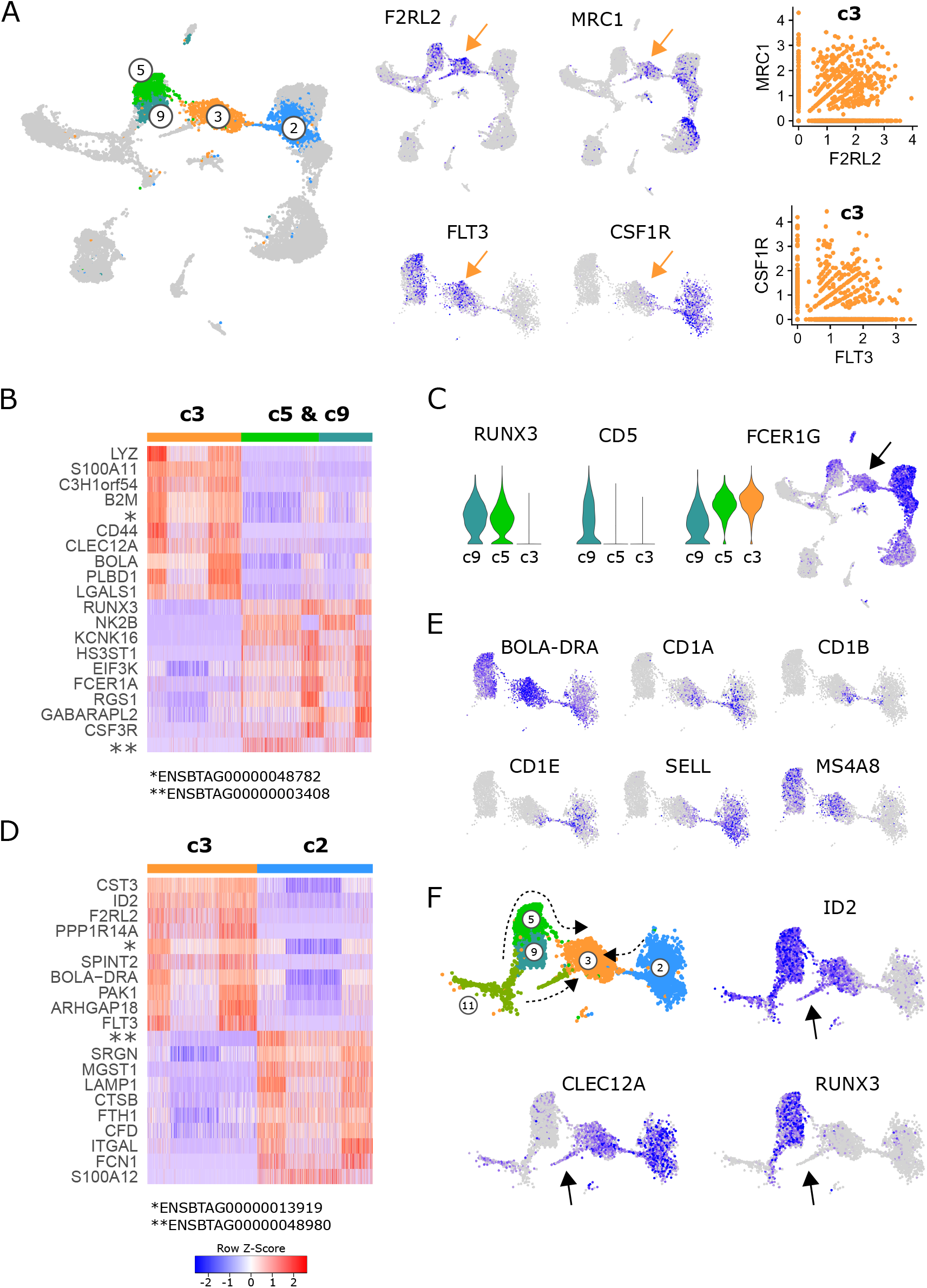
Dendritic cells and monocytes in cluster 3. (A) Expression of DC-associated and monocyte-associated genes in cluster 3. (B) Heatmap of the top 10 (p_adj) differentially expressed genes between cluster 3 and the cDC2 clusters c5 & c9. (C) Visualization of selected genes differentially expressed between c3, c5 and c9. Arrow indicates cluster 3. (D) Heatmap of the top 10 (p_adj) differentially expressed genes between cluster 3 and the monocytic cluster 2. (E) Visualization of selected genes in feature plots showing c2, c3, c5, and c9. (F) Proposed differentiation pathways of inflammatory cDC2 (c11→c9→c5→c3), moDC (c2→c3) and DC3 (c11→c3) indicated by dashed arrows. Arrows in feature plots indicate location of putative DC3 progenitors.

When compared against all other clusters, the most significant signature genes of cluster 3 included *PLBD1* (shared with cDC1), *CKB* and *VIM* **(Supplementary File 2)**. While *PLBD1* might be involved in the generation of lipid-based inflammatory mediators (44), *CKB* is implicated in immunometabolism of cells with high and fluctuating energy demands (45, 46), and vimentin (*VIM*) is described to be required for activation of the NLRP3 inflammasome (47).

When cluster 3 was compared with cDC2 (c5 & c9), the antibacterial and pro-inflammatory gene transcription of cluster 3 became even more apparent, with high expression of *LYZ* (lysozyme) and *S100A11* (an alarmin), alongside *XDH*, described to promote NLRP3 inflammasome activation (48). **(Figure 4B and Supplementary File 8)**. Among the most significant genes with lower expression in c3 compared to cDC2 (c5 & c9) were *RUNX3, NK2B, FCER1A* and *CSF3R*.

It is likely that cluster 3 represents inflammatory cDC2, which have recently been described in mice (29). While we could hardly detect any expression of *FCER1A*, we found high expression of *FCER1G* in putative inflammatory cDC2 (c3). Notably, *FCER1G* was also expressed in cluster 5 of resident cDC2, with expression increasing from c9 via c5 towards c3. With *RUNX3* and *CD5* expression absent from c3, this may indicate a differentiation pathway from c9 (RUNX3^+^CD5^+^FCER1G^dim^) via c5 (RUNX3^+^CD5^−^FCER1G^+^) towards c3 (RUNX3^−^CD5^−^FCER1G^high^) **(Figure 4C and F)**.

Compared to the monocyte cluster 2, c3 had higher expression of e.g. *CST3* and *ID2* **(Figure 4D)**, both predominantely expressed by DC clusters, and higher expression of a number of genes coding for MHC-II molecules **(Supplementary File 8)**. Genes with lower expression in c3 compared to monocytes (c2) coded for C-C motif chemokine 23 (*ENSBTAG00000048980*), a granule associated proteoglycan (*SRGN*) and CD107a (*LAMP1*), as well as cathepsin B (*CTSB*), numerous inflammation-related proteins, and surface molecules CD11a (*ITGAL*) and CD172a (*SIRPA*) **(Figure 4D and Supplementary File 8)**. Moreover, while all cells in cluster 3 expressed high levels of *BOLA-DRA*, mainly cells bridging between c3 and c2 were found to express *CD1A, CD1B* and *CD1E* **(Figure 4E)**, suggesting the presence of monocyte-derived DC (moDC) specialized in lipid antigen presentation in cluster 3. In line with their blood-borne monocyte-origin, these putative moDC expressed *SELL* (CD62L) which mediates LN entry via high endothelial venules.

Notably, while *MS4A8* was found to be highly expressed selectively in cDC2 (c5, c9) and putative inflammatory cDC2 (c3, left part), it appeared to be absent from putative moDC (c3, right part). The function of MSA48 is poorly characterized, but it was recently found to be highly upregulated in bovine pregnant endometrium (49) and in blood of Salmonella-infected pigs (50).

Alongside inflammatory DC originating from resident cDC2, and moDC originating from blood-borne monocytes, cells of the recently described distinct DC lineage DC3 (31, 32) may be present in cluster 3. Indeed, the observation that cycling (*PCNA* expressing) pre-DC (c11) branch towards cluster 3 **(Figure 2A and F)**, would support the presence of DC3 as a developmentally distinct linage within cluster 3 **(Figure 4F)**. In line with DC3 progenitors described in humans (32), this separate branch of DC progenitors expressed *CLEC12A* alongside *ID2*. Moreover it lacked detectable *RUNX3* expression (indicated by black arrows in **Figure 4F**). Certainly, more detailed analyses are warranted to untangle what seems to be transcriptomic co-clustering of inflammatory cDC2, monocyte-derived DC and DC3 in bovine mesenteric lymph node.

Cluster 15, another cluster positioned in-between DC and monocytes in the UMPA plot, did not show any specific expression of marker genes **(Supplementary File 2)**. Instead, these cells stood out by a low number of detected UMIs and a low number of detected genes, while the proportion of reads mapping to ribosomal and mitochondrial genes was comparable to other clusters (data not shown).

### Separate clustering of monocytes and macrophages

Like in humans, monocytes in blood of cattle are divided into classical and nonclasssical monocytes, and an intermediate subset, based on CD14 and CD16 expression (11, 21, 22). In bovine lymph nodes, *CSF1R* and *SIRPA* expressing monocytic cells appeared to predominantly express either *CD14* (c4, c14) or *FCGR3A* (CD16; c7, c10) **(Figure 5A)**.

**Figure 5.**
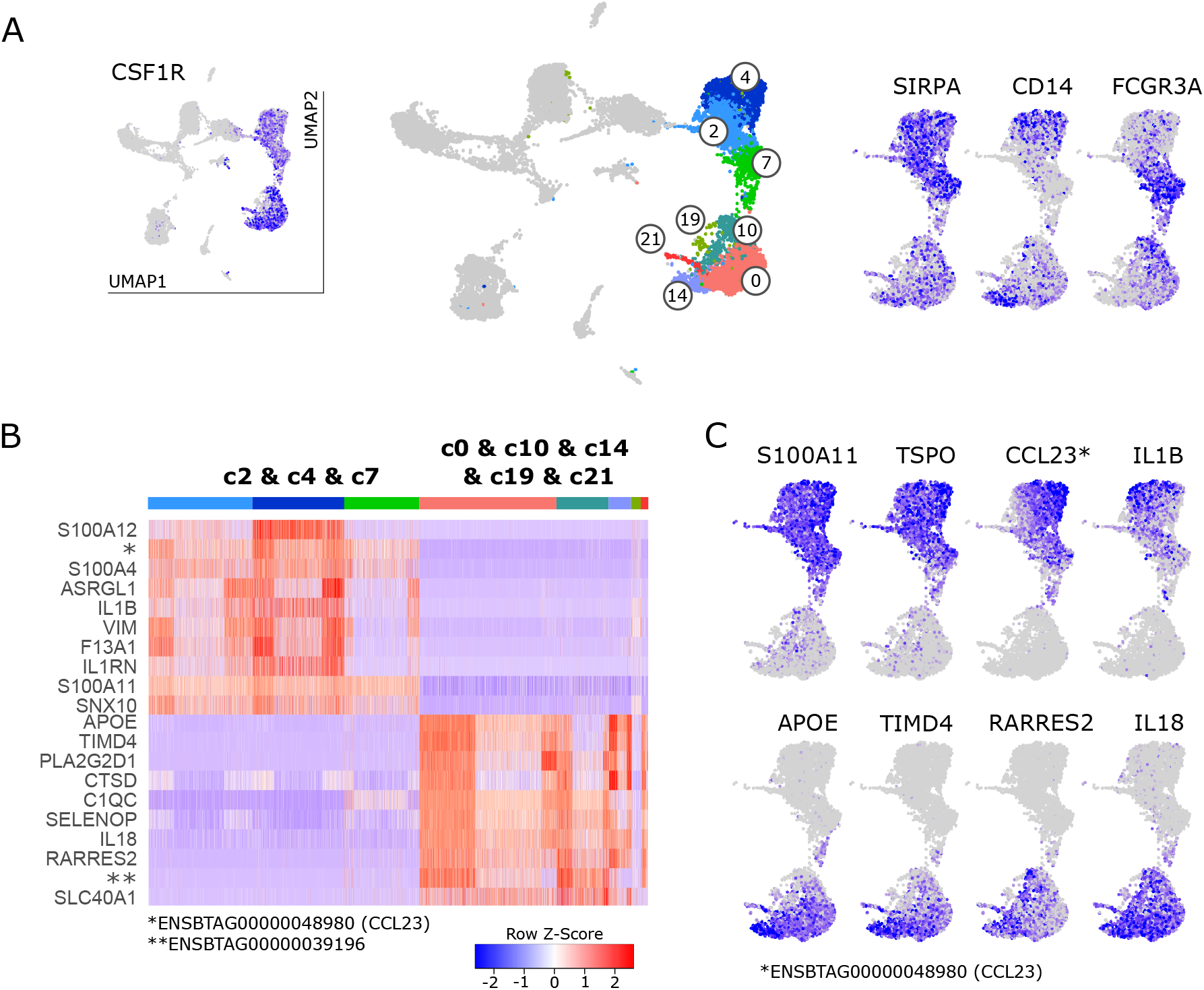
Monocytic cells. (A) Visualization of key genes that define monocytes and their subsets in blood of cattle. (B) Heatmap of the top 10 (p_adj) differentially expressed genes between monocyte clusters (c2 & c4 & c7) and macrophage clusters (c0 & c10 & c14 & c19 & c21). (C) Visualization of selected genes enriched in monocytes (top row) and macrophages (bottom row).

However, the pattern of *CD14*/*FCGR3A* expression did not reflect the observed division of monocytic clusters into two main islands in the UMAP projection: clusters 2 & 4 &7 and clusters 0 & 10 & 14 & 19 & 21. Differential gene expression between these two groups of clusters, revealed that one group (2&4&7) was clearly enriched in genes associated with classical monocytes found in blood of cattle (22) such as *S100A11, S100A12* and *VIM*, whereas the other group (0&10&14&19&21) was strongly enriched in transcripts for genes commonly associated with macrophages (*APOE, TIMD4, CD68*) **(Figure 5B and Supplementary File 9)**. Notably, putative monocytes exclusively expressed high levels of a gene recently annotated as CCL23 (*ENSBTAG00000048980*) and expressed high levels of *TSPO* and pro-inflammatory *IL1B*, whereas putative macrophages stood out by expressing *RARRES2* (Chemerin), *IL18*, and anti-inflammatory *PLA2G2D1* **(Figure 5B and C)**.

### Pro-and anti-inflammatory monocyte clusters

Monocyte clusters (c2, c4, c7) differed in the expression of genes related to inflammation and antigen presentation **(Figure 6 and Supplementary File 10)**. Top gene transcripts enriched in cluster 4 were reminiscent of pro-inflammatory classical monocytes in blood of cattle **(Figure 6A)**. When visualizing these genes in feature plots, expression gradients became apparent with highest expression levels for *S100A8, S100A12, VCAN*, and *DEFB7* in the top right corner of cluster 4 and tendentially higher expression of *IL1B* and *IL1RN* in the top left corner of cluster 4. Yet another expression pattern could be observed for *HIF1A* and *SLC2A3*, with decreasing expression from top to bottom of cluster 4 **(Figure 6B and Supplementary File 10)**. Cluster 2 was significantly enriched in transcripts related to antigen presentation such as *CD1E, BOLA-DRA, CD1A, CD74*, and *BOLA-DQA1* **(Figure 6A)**. Notably, expression of *CD1E, CD1A* and *CD1B* was mainly detected in cells of cluster 2 that seemed to bridge towards dendritic cells (i.e. cluster 3, see also **Figure 4E**). Moreover, several C-type lectin receptors, most prominently *CLEC6A*, showed increased expression in cells of cluster 2 **(Figure 6C and Supplementary File 10)**. Cluster 7 was significantly enriched in gene transcripts associated with nonclassical monocytes in blood of cattle, such as *C1QA, C1QB, C1QC, FCGR3A*, and *CX3CR1*. Notably, expression of *MS4A7* was shared between clusters 2 and 7, and was almost absent from pro-inflammatory cluster 4 **(Figure 6D and Supplementary File 10)**.

**Figure 6.**
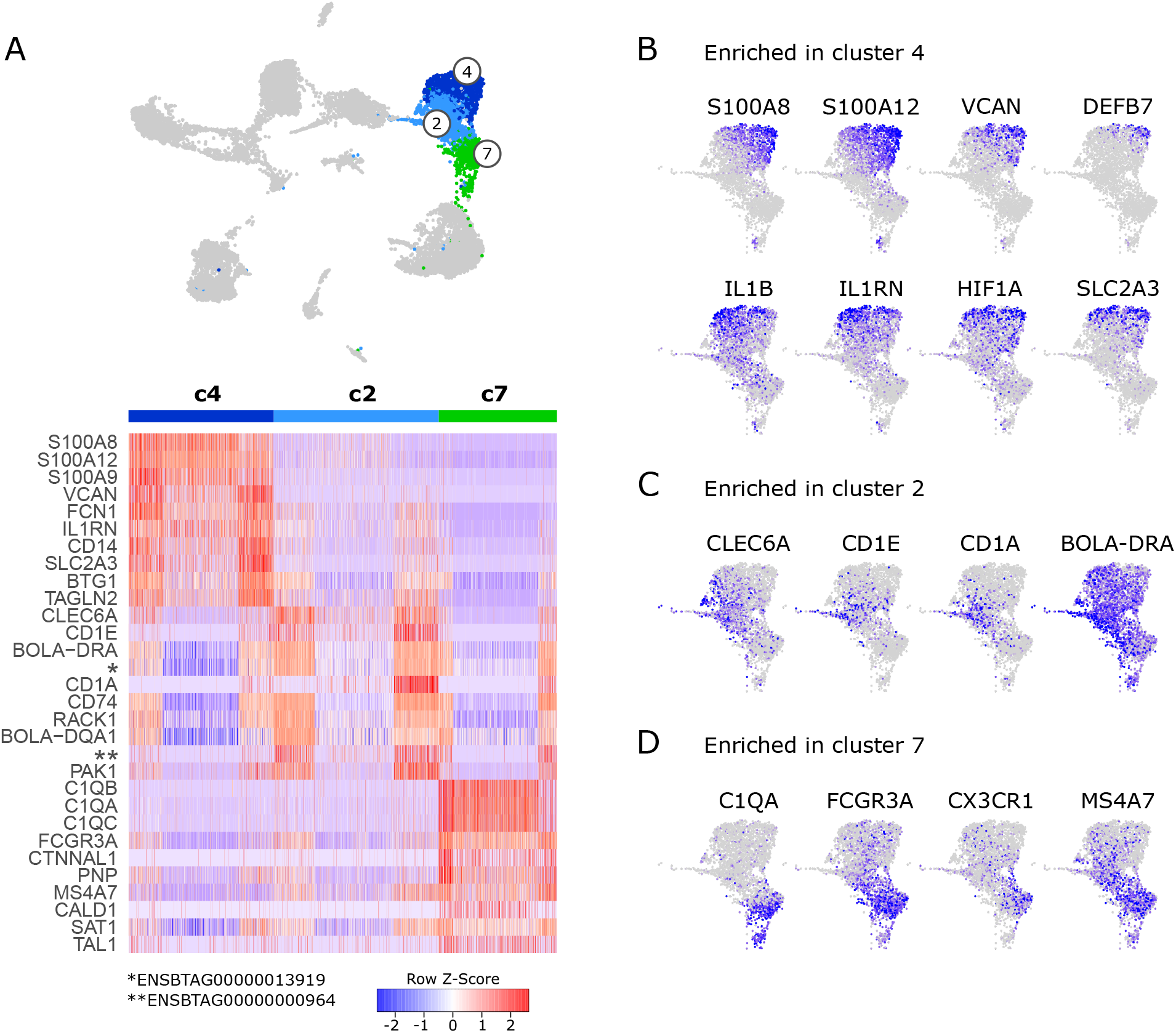
Monocytes. (A) Heatmap of the top 10 (p_adj) differentially expressed genes between monocyte clusters (c2, c4, c7). (B-D) Visualization of selected genes enriched in cluster 4 (B), cluster 2 (C), and cluster 7 (D).

### Macrophage clusters

The majority of macrophages clustered together as cluster 0. While no genes were found to be exclusively expressed in c0, the clusters sourrounding c0 showed a more pronounced differential gene expression signature **(Figure 7A)**. Notably, cells in cluster 10 appeared to uniquely express *CD5L, SIGLEC1* (CD169) and *CD163* **(Figure 7A and B)**. Also genes encoding the phagocytic receptors *MRC1* (CD206) and *CLEC4F* (51) were predominantly expressed in c10, as was *HMOX1*, encoding a heme-degrading enzyme with anti-inflammatory effects (52). Moreover, cells in c10 expressed significantly increased levels of regakine-1 (*ENSBTAG00000010155*), a CC-chemokine found to be constitutively present at high concentrations in bovine plasma and to attract neutrophils and lymphocytes *in vitro* (53).

**Figure 7.**
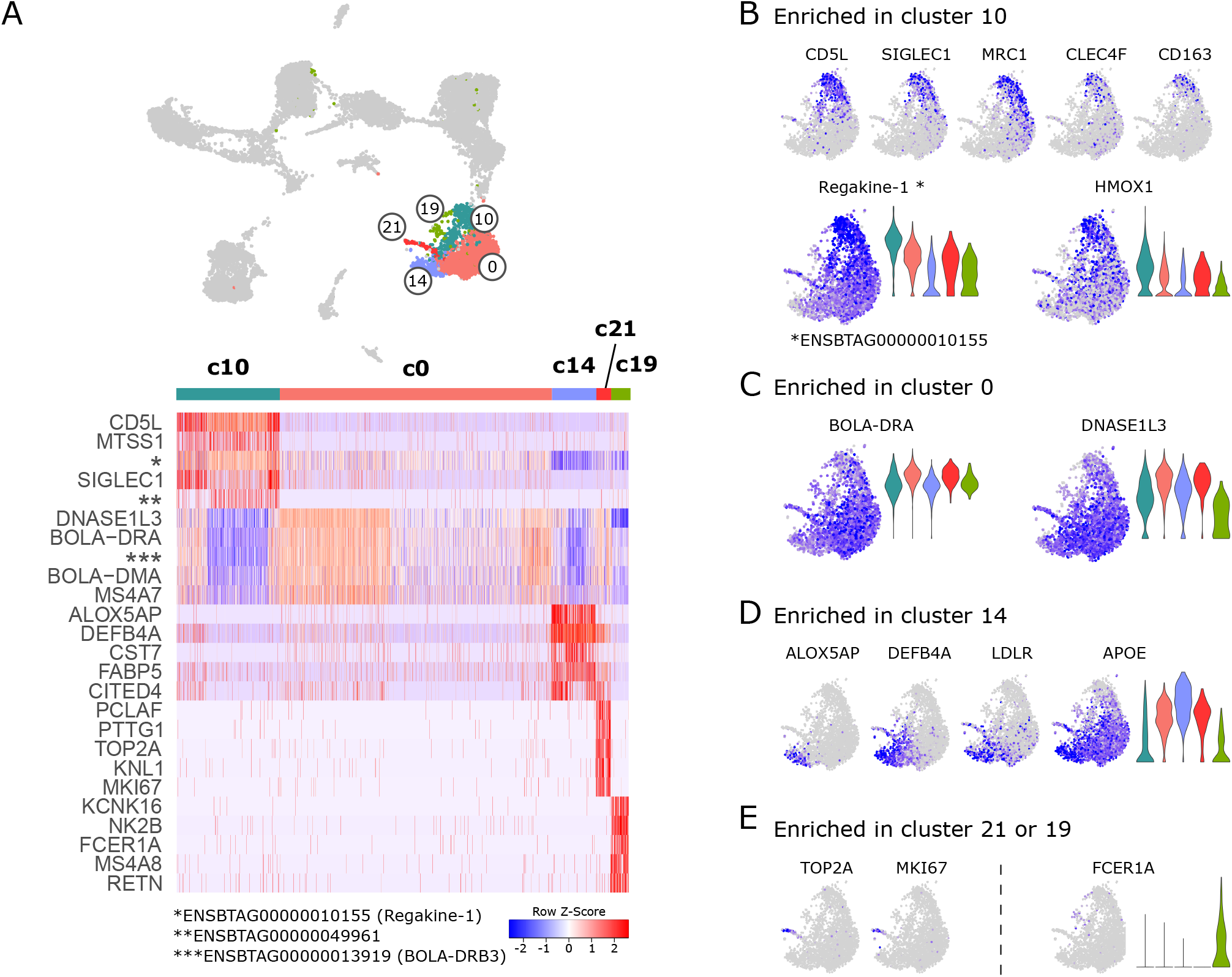
Macrophages. (A) Heatmap of the top 10 (p_adj) differentially expressed genes between macrophage clusters. (B-E) Visualization of selected genes enriched in cluster 10 (B), cluster 0 (C), cluster 14 (D), and in clusters 21 and 19 (E).

Similar to monocytes in cluster 2, macrophages in cluster 0 expressed higher levels of genes related to antigen presentation (*BOLA-DRA, BOLA-DMA, BOLA-DQA1, BLA-DQB, CD74*) **(Figure 7C and Supplementary File 11)**. High expression levels of *DNASE1L3, MERTK* and *AXL* in cluster 0, point towards a prominent role of these macrophages in efferocytosis.

Together with pro-inflammatory monocytes, cells in cluster 14 uniquely expressed *ALOX5AP* and *DEFB4A*. While ALOX5AP is crucial for leukotriene biosynthesis, DEFB4A is described to have both antimicrobial and chemotactic functions (54). Other genes predominantly detected in cluster 14 encoded cystatin F (*CST7*), reported to regulate proteolytic activity during monocyte-to-macrophage differentiation (55), the receptor for low-density lipoprotein (*LDLR*), and the fatty-acid binding protein 5 (*FABP5*). Expression of FABP5 is in line with the idea that c14 represents pro-inflammatory macrophages, as this fatty-acid binding protein was shown to limit the anti-inflammatory response of murine macrophages (56).

Cells in cluster 21 stood out by high expression of proliferation-associated genes such as *TOP2A* and *MKI67*. When c21 was compared against the proliferating DC clusters 11 & 12, the clear macrophage identity (e.g. *CD68, CSF1R*) of c21 became apparent (data not shown). The presence of proliferating macrophages in the current dataset may suggest that some macrophages in lymph node of cattle are replaced by self-renewal, as reported for bona fide tissue-resident macrophages (25).

Cells in cluster 19 shared a number of genes with cDC2. Cells of this cluster were also located together with cDC2 and monocytes in the UMAP projection. Due to the small size of cluster 19 and the widespread positioning of its cells, we concluded that c19 might represent an artefact and should be interpreted with caution.

### Genes of interest

While differential expression testing is restricted to and limited by cluster definitions, visualization of genes in feature plots provides unbiased and cluster-independent information, which is highly valuable to reveal sub-clustering and expression patterns that are hidden in the cluster analysis. So in addition to the differential expression testing described above, we examined genes associated with important immunological functions, such as pattern recognition, adhesion, migration, and antigen presentation, as well as genes coding for cytokines and their receptors and other immuno-relevant molecule classes, such as solute carriers, tetraspanins, semaphorins, metalloproteinases and purinergic receptors.

Very interesting patterns of gene expression became apparent which cannot all be described in detail here. Selected feature plots are shown in **Figure 8**. The complete collection of feature plots is given in **Supplementary File 12**. When interpreting these feature plots, the reader should keep in mind that dropout in single-cell data might contribute to low detection and apparent lack of expression.

**Figure 8.**
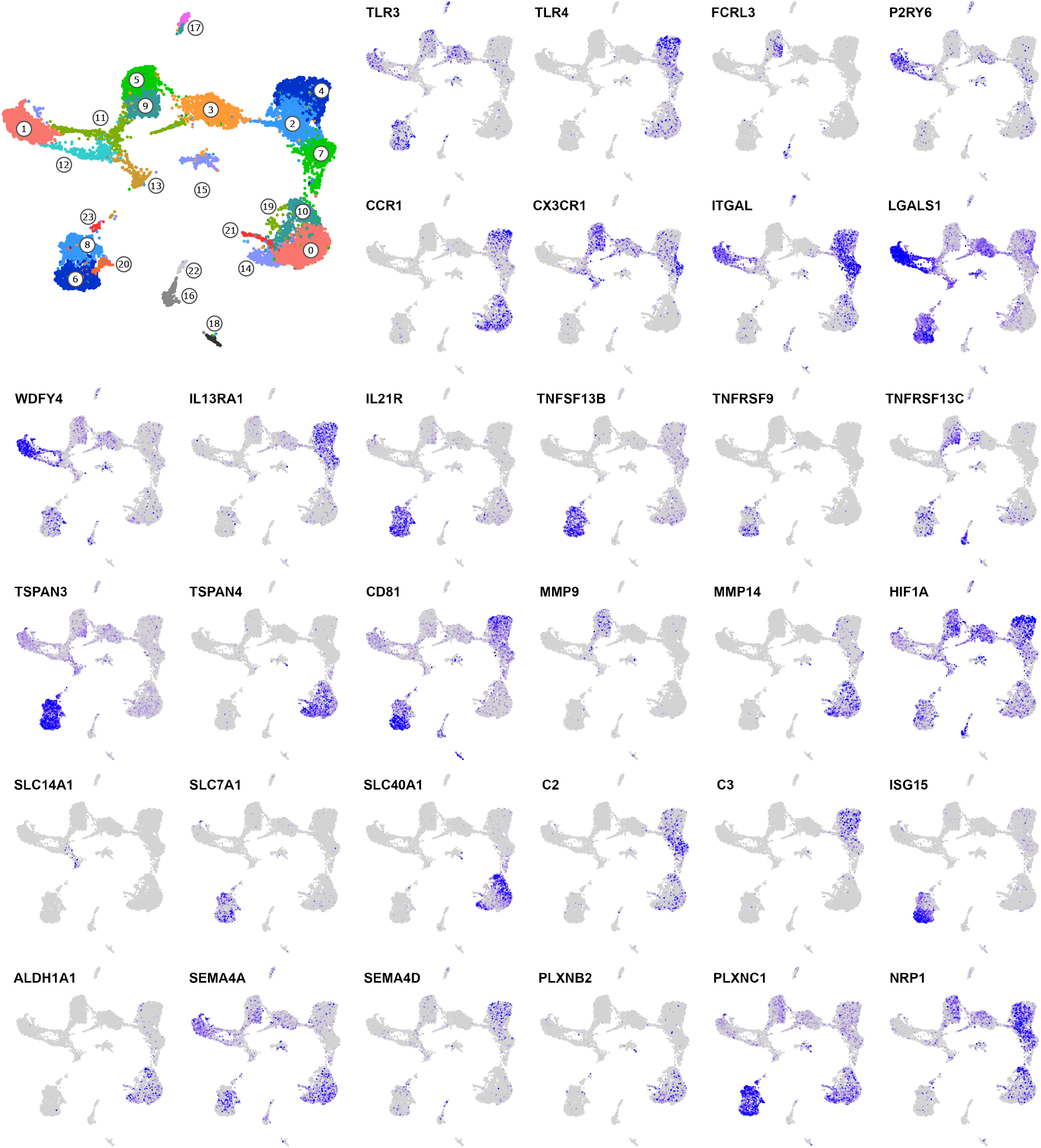
Genes of interest. The complete collection of feature plots is given in Supplementary File 12 and includes the following categories: 1) Pattern recognition receptors, 2) Fc receptors, 3) Purinergic receptors, 4) Chemokines, 5) Chemokine receptors, 6) Integrins, 7) Galectins, 8) Antigen presentation, 9) T-cell modulation, 10) Interleukins, 11) Interleukin receptors, 12) TNF superfamily, 13) TNF receptor superfamily, 14) Tetraspanins, 15) Metalloproteinases, 16) Metabolism (misc.), 17) Glycolysis, 18) Solute carriers, 19) Complement system, 20) Interferon-associated, 21) Retinoic-acid production and signaling, 22) Semaphorins and receptors.

Transcripts for **pattern-recognition-receptor** genes were predominantly detected in monocytic cells. Apart from for example *TLR3* transcripts, which were mainly detected in dendritic cells, and *CLEC9A* transcripts, which were exclusively detected in resident cDC1 (c1 and part of c11/c12), who also expressed high levels of *CLEC12A* alongside cluster 3 (inflammatory cDC2) and monocytes. Interestingly, *NLRP3* expression appeared to be limited to monocytic cells and cDC2 (including c3), while *NLRP1* was also highly expressed in cDC1 (c1).

Looking at **Fc receptors**, we found *FCER2* to be expressed predominantly in monocytes and in cluster 3 (inflammatory cDC2), as well as in putative migratory cDC2 (c8). Notably, while *FCER1G* expression was detected in *CD5*^−^ cDC2 (c5), but not in *CD5*^+^ cDC2 (c9), expression of *FCRL3* and *FCRL4* followed the opposite pattern, with clear transcript enrichment in CD5^+^ cDC2 (c9).

Among transcripts encoding **purinergic receptors**, involved in immune regulation by sensing extracellular nucleotides (57), *P2RY6* transcripts were clearly enriched in resident cDC1 (c1). In line with our previous results obtained with blood-derived cells of cattle (11), *P2RY1* and *P2RY10* transcripts were primarily detected in monocytic and dendritic cells, respectively.

Also **chemokines and chemokine receptors** showed clear cluster-specific expression patterns. Most strikingly, *CCR7*-expressing DC could be divided according to *CCL17* / *CCL22* expression (c8, putative migratory cDC2) and *CXCL9* / *CXCL10* expression (c6, putative migratory cDC1). Apart from c6, *CXCL9* and *CXCL10* were also highly expressed in what appears to be a subcluster of c1 (resident cDC1), presumably containing activated resident cDC1. Notably, *CX3CR1* was mainly expressed by DC progenitors, cDC2 (including c3) and monocytes. Apart from *CX3CR1* and *CCR5*, we could hardly detect any transcripts coding for chemokines or chemokine receptors in progenitor cells (c11, c12, c13), cDC2 (c5, c9) and cluster 3. Remarkable is also the high expression of *CCR1* and *CXCR4* in pro-inflammatory monocytes (c4). However, while *CXCR4* expression decreased towards anti-inflammatory monocytes and macrophages, *CCR1* expression was again detected at high levels throughout macrophage clusters. Moreover, expression of the follicle-homing receptor *CXCR5* was exclusively detected in putative migratory cDC2, suggesting that these cells locate close to follicles to interact with B- and T-cells, as described for the murine system (58).

Among the investigated **integrins**, *ITGAL* (CD11a) stood out by exclusive high expression in resident cDC1 (c1) and monocytes (c4, c2, c7). And transcripts for *ITGAV* (CD51) were almost exclusively detected in pro-inflammatory macrophages (c14).

In addition, **galectins** showed specific expression patterns. Most notably Galectin-1 (*LGALS1*), which was highly expressed in both resident and migratory cDC1 (c1, c6) and in inflammatory and migratory cDC2 (c3, c8). Galectin-1 expression in DC is reported to induce IL-27 expressing regulatory DC (59). Transcripts for *LGALS3* and *LGALS9* were enriched in macrophages and monocytes, respectively.

Moreover, genes related to **antigen presentation and T-cell modulation** showed interesting cluster-specific patterns. Notably, cluster 6 (putative cDC1 within migratory DC), clearly showed the highest levels of *BOLA, BOLA-NC1* and *IDO1*, suggesting that these cells interact with and regulate CD8 T cells. Resident cDC1 (c1) were clearly enriched for *WDFY4* transcripts, presumably indicating their potential for cross presentation. Notably, while *CD1A, CD1B* and *CD1E* were exclusively expressed by putative moDC (bridging between c2 and c3), *CD1D* was also highly expressed by resident cDC1 (c1) and to a lesser extent by cDC2 (c3, c5, c9).

Looking at **interleukins**, we found that *IL1B* was expressed mainly by monocytic cells and cDC2 (including c3), while the gene for the IL1 receptor antagonist (*IL1RN*) was mainly detected in pro-inflammatory monocytes (c4). Transcripts for IL-18 (*IL18*) showed a prominent and specific expression in macrophages. Notably, subclusters of migratory DC clearly differed in their expression of *IL15* (c8, putative migratory cDC2) and *EBI3* (IL-27B) (c6, putative migratory cDC1). Among **interleukin receptor** genes, *IL21R* and *IL13RA1* stood out by their exclusive expression in migratory DC and monocytes, respectively.

Genes encoding members of the **TNF- and TNF-receptor superfamily** were also differentially expressed, with *TNFSF13* (APRIL) and *TNFSF10* (TRAIL) being detected primarily in pro- and anti-inflammatory monocytes (c4 vs. c7), respectively. Notably, transcripts for TNF-α (*TNF*) were only poorly detected, but expression of the TNF receptor genes *TNFRSF1A* (TNFR1) and *TNFRSF1B* (TNFR2) was detected at high levels in monocytic cells (*TNFRSF1A/B*) and migratory DC (*TNFRSF1B*). Moreover, while *TNFSF13B* (BAFF) was highly and almost exclusively expressed in all migratory DC, transcripts of the TNF receptor genes *TNFRSF9* and *TNFRSF13C* were enriched in putative migratory cDC1 (c6) and CD5 expressing resident cDC2 (c9) / putative migratory cDC2 (c8), respectively.

Also **tetraspanins** showed pronounced cell-type specific expression, for example the macrophage-restricted expression of *TSPAN4*, recently shown to interact with Histamin H4 receptor (60), or the high transcription of *TSPAN3* in migratory DC, among which migratory cDC1 (c6) were also enriched in *CD151* (TSPAN24) and *CD81* (TSPAN28) transcripts. Other tetraspanin genes highly expressed in migratory DC included *TSPAN13, TSPAN17*, and *TSPAN33*, the two latter of which are known to interact with ADAM10, a metalloproteinase mediating ectodomain shedding (61). While *ADAM10* transcripts were detected in all clusters with higher levels in monocytic cells, other **metalloproteinases** showed more specific patterns of expression. Transcripts for ADAM8 and ADAM9 were enriched in migratory DC (c6, c8), and transcripts for MMP9 and MMP14 were almost exclusively detected in resident cDC2 (c5, c9) and macrophages (c0, c10, c14, c21), respectively.

Among genes involved in **metabolism** and previously reported to be overexpressed in cM compared to ncM in bovine blood (22), *KHK* and *SORD* (fructose metabolism) stood out by selectively higher expression in monocytic clusters and in resident cDC2 (c5, c9), respectively. Transcripts involved in **glycolysis** showed surprisingly heterogeneous expression patterns, but were mostly detected in monocytes and cDC2 (including c3), consistent with the highest transcript levels of *HIF1A* in these clusters. In line with the pro-inflammatory signature of monocytes in c4, these cells transcribed the highest levels of *SLC2A3*, a high-affinity glucose transporter.

Visualization of SLC (**solute carrier**) gene expression revealed interesting patterns that are also reported for humans, like high *SLCO2B1* expression in macrophages (62) or high *SLCO5A1* expression in mature dendritic cells (63). Notably, transcripts for the iron transporter *SLC40A1* were only detected in certain cluster-independent regions within macrophages, and the urea transporter *SLC14A1* was exclusively detected in putative early DC progenitors (c13). Moreover we found that migratory DC exclusively expressed high levels of *SLC7A1*, which was recently described as a cellular receptor for bovine leukemia virus (64). Infection of migratory DC by bovine leukemia virus may have important implications for pathogenesis of the disease. Notably, bovine leukemia virus has recently also gained attention as a potential causative agent in human breast cancer (65).

Interesting expression patterns were also apparent for genes related to the **complement system**. Transcripts for C1Q, the sensory component initiating the formation of the C1 complex (C1Q+C1R+C1S), were exclusively expressed in macrophages and anti-inflammatory monocytes. The latter (c7) also expressed the highest levels of *C2*, while transcripts for C3 were exclusively detected in rather pro-inflammatory monocytes (c4, c2). Conversely, *C3AR1* transcription appeared to be selectively absent from pro-inflammatory monocytes (c4). Notably, detection of *CD55* transcripts was limited to monocytes. Complement factor D (*CFD*) and *CFP*, involved in the alternative pathway of complement activation, were also detected in dendritic cells, *CFD* most prominently in non-proliferating progenitors (c13), putative migratory cDC2 (c8), as well as in CD5^−^ resident cDC2 (c5) and cluster 3 (inflammatory cDC2).

Among **interferon-associated** genes, *ISG15* and *IFI6* stood out by their almost exclusive and high expression in putative migratory cDC1 (c6) and macrophages, respectively.

**Retinoic acid** (RA) is reported to imprint gut homing in T cells (66, 67) and – as recently reviewed – to control IgA switch in B cells (68). As reported in these studies, mucosal subsets of myeloid cells appear to be specialized to produce RA, reflected by their unique expression of required enzymes. Notably, in our dataset we found high expression of *ALDH1A1* exclusively in macrophages, and some *ALDH1A2* expression in cells assigned to resident cDC1 and migratory DC clusters. We also detected CD103 (*ITGAE*) expression in DC, which is regarded as a marker for “RA-DC” in mice, alongside expression of retinoic acid receptors (*RARA, RARB, RARG, RXRA, RXRB*). Notably, the RA-responsive genes *RARRES1* and *RARRES2* (Chemerin) were predominantly expressed in macrophages, the latter also in migratory cDC1.

Finally, genes for **semaphorins** (*SEMA4A, SEMA4D, SEMA4F, SEMA7A*) and their receptors (plexins such as *PLXNB2* and *PLXNC1* and neuropilins such as *NRP1*) showed unique expression patterns across dendritic and monocytic clusters. Notably, semaphorin signaling is regarded as highly conserved across species (69) and its regulatory roles in innate immunity are only beginning to be elucidated (70).

### Trajectories and sources of MPS heterogeneity in bovine mesenteric lymph node

Our scRNA-seq analysis of mononuclear phagocytes in bovine mesenteric lymph nodes revealed clusters of resident DC (progenitors, cDC1, cDC2, pDC), as well as migratory DC (putative cDC1 and cDC2), and a cluster of inflammatory cDC2 containing moDC and potentially DC3 **(Figure 9A)**. Monocytic cells could be clearly separated into monocytes and macrophages, both clustering according to pro-and anti-inflammatory gene expression, and included a cluster of proliferating macrophages, presumably giving rise to bona fide lymph-node resident macrophages. As illustrated in **Figure 9B**, heterogeneity of mononuclear phagocytes in bovine lymph nodes may originate from blood-borne progenitors (S1) that differentiate into resident cDC1 (T1a and T1b), resident cDC2 (T2) and resident DC3 (T3). Highly activated migratory DC (S2) may enter via afferent lymph. Furthermore, cDC2 may differentiate into inflammatory cDC2 (T4). Monocytic cells may originate from cM that enter the lymph node (S3) and differentiate into antigen-presenting moDC (T5) or via antigen-presenting monoctyes (resembling intM in blood) towards increasingly anti-inflammatory monocytes (resembling ncM in blood) and further into macrophages (T6). Macrophages may aquire anti-inflammatory or pro-inflammatory gene expression depening on the niche they occupy. Alternatively, they may aquire an increasingly pro-inflammatory gene expression over time (T7). Lastly, self-renewing tissue macrophages (S4) may contribute to the pool of macrophages (T8), displaying a transcriptome indistinguishable from terminally differentiated monocyte-derived macrophages.

**Figure 9.**
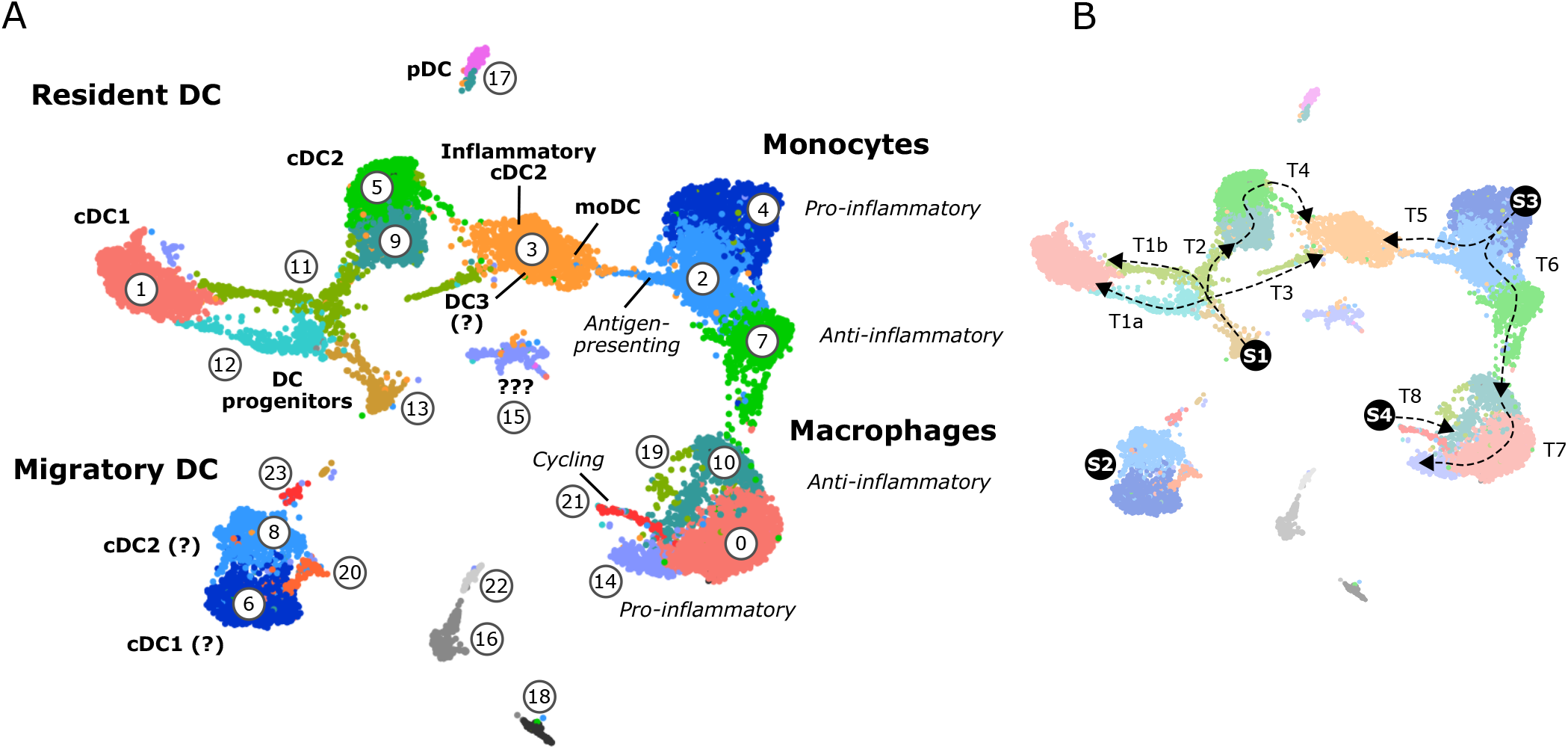
Heterogeneity of mononuclear phagocytes in bovine mesenteric lymph node. (A) Overview of cluster assignment. (B) Proposed source populations (S1-S4) and proposed differentiation trajectories (T1-T8).

## DISCUSSION

In the present study, we have applied 10x Genomics and Illumina sequencing to decipher the single-cell transcriptome of mononuclear phagocytes in bovine mesenteric lymph nodes.

One particularly interesting aspect of the present dataset is the collection of single-cell transcriptomes from migratory DC subsets, enabling an unbiased view on *in-vivo* activated DC and their profound transcriptional reprogramming. We identified two subclusters of migratory DC that clearly differ in their expression of chemokine genes and likely contain cDC1 (c6; *CXCL9, CXCL10*) and cDC2 (c8; *CCL17, CCL22*). High expression of interferon-inducible genes *CXCL9* and *CXCL10* might indicate that migratory cDC1 occupy a niche where they interact with CXCR3-expressing Th1 and CD8 T cells. Indeed, high expression of *BOLA* (MHC-I) and *B2M* would support a specialization towards CD8-T-cell interaction. Notably, *CXCR3* transcripts were found in resident cDC1 alongside *CXCL9* and *CXCL10* transcripts, suggesting a common niche of resident and migratory cDC1 for promoting antiviral responses. A recent study performed in mice suggested that CXCL9 and CXCL10 are produced by spatially distinct DC subsets, creating distinct microenvironments and favoring distinct effector and memory T-cell fates (71). In our dataset, most migratory DC contained transcripts for both CXCL9 and CXCL10, making compartmentalization purely based on differential expression of these two chemokines rather unlikely, at least in the bovine system.

Dominant expression of *CCL17* and *CCL22* detected in putative migratory cDC2 would suggest that these cells are specialized in attracting CCR4-expressing Th2/Th17 cells as well as Tregs (43), presumably creating a niche supporting survival and proliferation of these cells by IL-15 expression. Moreover, some of these putative migratory cDC2 were enriched in *CXCR5* transcripts, suggesting that they locate close to B-cell follicles to support follicular T-helper cell (Thf) differentiation (2). In this regard, production of soluble CD25 by DC has been proposed as a means to support Thf differentiation (72). Transcripts for CD25 (*IL2RA*) were poorly detected in the current dataset, but were primarily found in clusters associated with cDC2, including cluster 8. We have previously reported pronounced upregulation of surface CD25 expression after 4-hour *in-vitro* stimulation of bovine DC subsets (33), making expression of membrane-bound and/or soluble CD25 by activated DC a likely mechanism to control Thf differentiation also in bovine lymph nodes.

Apart from activation in the periphery, some *CCR7*^+^ DC may also have differentiated from resident DC that were activated in the lymph node and have upregulated *CCR7* expression in order to migrate towards T-cell zones, as recently described for mice (4). In fact, a few cells adjacent to resident cDC1 also expressed *CCR7*, and displayed a more activated transcriptome. Moreover, both tolerogenic and immunogenic DC may be present in the migratory DC clusters, as molecular changes have been shown to be highly similar for both activation states (73). Along this line, recently described mregDC (74) may be included as well, given the prominent expression of regulatory genes such as *CD274* (PDL-1), *PDCD1LG2* (PDL-2), *FAS*, and *SOCS2* in *CCR7*-expressing DC.

The recent discovery of DC3 as a separate DC lineage (31, 32) prompted us to watch out for these cells in the current dataset. Looking at the branching pattern of DC progenitors, it is conceivable that some cells in cluster 3 constitute DC3, however we could not find a clear separation from putative inflammatory cDC2 and monocyte-derived DC. Certainly, the seemingly convergent differentiation pathways of several cell types might indicate crucial roles of cells with this transcriptomic makeup – it remains to be determined if these inflammatory cells act in different niches of the lymph node with nuanced differences in their specialization, or if there is pronounced redundancy in the system because the tasks they fulfill are crucial for proper functioning of the immune system and thus survival.

The high proportion of monocytic cells in bovine mesenteric lymph nodes, and reportedly also in murine and human lymph nodes (6), raises several questions regarding their functional roles, their origins, and their fates. The idea that dendritic and monocytic cells cooperate in the lymph node to optimize adaptive immune responses, is supported by recent findings in mice, where inflammatory monocytes were shown to enter the lymph node from blood via high endothelial venules (HEV) and accumulate in the T-cell zone, where they provide polarizing cytokines to optimize effector T-cell differentiation (4). In line with this, we found monocytic cells in bovine mesLN to express high levels of *SELL* (CD62L) alongside *CXCL16*, attracting activated T cells (75), and transcripts for T-cell modulating cytokines, such as IL-16, IL-12B/IL-23A and IL-27/ IL-27B (*EBI3*). Classical monocytes were also described to travel to lymph nodes via tissue-draining lymph, both under inflammatory and steady-state conditions (76). Accordingly, murine moDC were reported to be capable of CCR7 upregulation *in vivo* (77). It is unclear if bovine monocytes can upregulate CCR7 *in vivo*. At least *in-vitro* activation with TLR-ligands did not lead to an increase of CCR7 transcripts and protein expression in bovine cM (33). Moreover, in the present study we did not detect any CCR7 transcripts in lymph-node derived monocytic cells.

Our recent scRNA-seq analyses of bovine blood monocyte subsets support the idea of continuous differentiation from cM via intM to ncM (22). It remains to be determined if classical pro-inflammatory monocytes that enter the lymph node can differentiate to cells that resemble intM and ncM found in peripheral blood of cattle, or if intM and ncM differentiated in blood can enter lymph nodes themselves. The almost complete absence of CD62L expression (mRNA and protein) from bovine intM and ncM in blood (22), would argue against them having access via HEV, or at least indicates differential regulation of LN entry.

The biology of tissue-resident macrophages is described to be shaped by four different factors: i) origin (embryonic vs. monocyte-derived), ii) tissue-specific environment (e.g. lung vs. liver), iii) inflammatory environment, and iv) time spent in the tissue (78). Consistent with described functions of macrophages, our data suggests a division of labor between rather anti-inflammatory macrophages engaging in efferocytosis, and macrophages with antibacterial activity and a rather pro-inflammatory profile. Furthermore, the detection of cycling macrophages in the present dataset clearly supports the hypothesis that self-renewing macrophages of embryonic origin are present in bovine lymph nodes. The fact that these bona fide macrophages don’t segregate as a separate cluster in our dataset, is in line with the idea that monocyte-derived macrophages, educated by niche-specific signals, acquire a very similar and thus undistinguishable transcriptome over time. Expression of *TIMD4* has been associated with long-term residence under steady-state conditions – thus being upregulated on monocyte-derived macrophages over time (25). In the current dataset, the continuous increase of *TIMD4* expression across macrophage clusters may therefore indicate a differentiation path for monocyte-derived macrophages. The expression of *APOE* followed the same pattern, suggesting that also *APOE* might serve as a time-dependent marker for macrophage differentiation.

It has been suggested that differentiation of monocytes to tissue macrophages can be split into two phases (78): a rapid differentiation that would instruct cells to stay in the tissue niche (“stay-here” signals), and a second phase, where monocyte-derived macrophages would adapt to their environment integrating information on tissue type and inflammatory status (“learn-this” signals). It is intriguing to speculate that this first rapid phase of differentiation is visible in the UMAP projection of our dataset, where monocytes and macrophages are connected with a narrow bridge of cells. The low frequency of transitional cells would be in line with rapid differentiation. Similarly, differentiation towards moDC appears to be a rapid process, leading to a narrow cellular bridge between monocytes and cluster 3 in our dataset.

Further macrophage heterogeneity is introduced by niches within the same tissue and stromal and immune cells present therein. For the lymph node, several niche-specific subsets of macrophages have been described (79), such as subcapsular and medullary sinus macrophages, capturing lymph-borne antigens (80, 81), macrophages in the lymph node parenchyma such as in germinal centers and medullary cords, and recently described efferocytotic T-cell zone macrophages (82). Our data reveals cluster-specific heterogeneity in the expression of chemokine receptor genes such as *CCR1, CCR5, CXCR4* and *CX3CR1*, presumably guiding monocytic cells to their “niche of residence”, where further differentiation may also depend on available space in that niche (83).

One limitation of the current study is that pDC were only detected in a very small cluster (c17). In fact, flow cytometric analyses suggest that the frequency of pDC in bovine mesLN is considerably higher than suggested by our scRNA-seq dataset (unpublished data). With the gating strategy employed to sort mononuclear phagocytes in the present study we aimed to reduce contamination with lymphocytes, but at the same time we excluded most pDC that express comparatively low levels of Flt3. A future study will have to address pDC in the lymph node and investigate for example if pDC subsets are present in a special cDC2-like activation state, as described for human and murine transitional DC (18, 27, 84–87). An indication for these transitional DC in the current dataset may be that a small subset of cells spatially clustering with pDC (c17) got assigned to a cDC2 cluster (c9).

With the present study we provide the first in-depth single-cell analysis of the mononuclear phagocyte compartment in bovine lymph nodes. Trajectories of differentiation became apparent that may well reflect general principles of MPS dynamics in lymph nodes across species. Some of them previously reported, such as the differentiation of resident DC from blood-borne progenitors in the lymph node (88, 89), others less well understood, such as the origin and differentiation pathways of monocytic cells and bona fide macrophages in secondary lymphoid tissue, or the seemingly convergent functions of inflammatory DC subsets (including DC3) and monocyte-derived DC.

The hypotheses generated in this manuscript are an important contribution towards a better understanding of the mononuclear phagocyte compartment in lymph nodes, especially when acknowledging that basic DC and monocyte biology appears to be largely conserved across mammalian species. Spatial analyses performed in future studies should help to define niche-specific transcriptomes of macrophages and to characterize microenvironments where dendritic cells, monocytes and T-cells interact to shape adaptive immune responses – fundamental insights into MPS biology that will benefit human and veterinary medicine alike.

## Supporting information

Suppl_1_CellRanger_Clustree

Suppl_2_complete_dataset

Suppl_3_resDC

Suppl_4_c5vsc9

Suppl_5_keyGenes

Suppl_6_migDCresDC

Suppl_7_migDC

Suppl_9_monMac

Suppl_10_mon

Suppl_11_Mac

Suppl_12_GOI

## ACKNOWLEDGEMENTS

We thank Corinne Hug (IVI) for producing recombinant bovine Flt3L, Stefan Müller (FCCS, University of Bern) for sorting, and Pamela Nicholson, Catia Coito and Tosso Leeb from the NGS Platform of the University of Bern for single-cell RNA sequencing.

## AUTHOR CONTRIBUTIONS

GTB performed laboratory work, analyzed data, and wrote a first draft of the manuscript. SCT performed data analysis, prepared the figures, and wrote the final manuscript. MCK and RB performed bioinformatic analyses. SCT and AS designed and supervised the overall project. All the authors reviewed the manuscript.

