## Supplementary material for "Single-cell transcriptomics reveals striking heterogeneity and functional organization of dendritic and monocytic cells in the bovine mesenteric lymph node": Suppl_1_CellRanger_Clustree

### Supplementary File 1A



#### MLN2306

**The analysis detected some issues. Details »**

| Alert | Value | Detail |
|---|---|---|
| ⚠ **Low Fraction Valid UMIs** | 69.4% | Ideal > 75%. This may indicate a quality issue with the Illumina R2 read for Single Cell 3' v1 or the R1 read for Single Cell 3' v2/v3 and Single Cell 5'. Application performance may be affected. |

##### Estimated Number of Cells

6,937

| Mean Reads per Cell | Median Genes per Cell |
|---|---|
| 120,798 | 898 |

##### Sequencing

| | |
|---|---|
| Number of Reads | 837,977,792 |
| Valid Barcodes | 97.1% |
| Sequencing Saturation | 61.7% |
| Q30 Bases in Barcode | 96.5% |
| Q30 Bases in RNA Read | 94.1% |
| Q30 Bases in UMI | 96.4% |

##### Mapping

| | |
|---|---|
| Reads Mapped to Genome | 95.7% |
| Reads Mapped Confidently to Genome | 88.5% |
| Reads Mapped Confidently to Intergenic Regions | 11.5% |
| Reads Mapped Confidently to Intronic Regions | 19.5% |
| Reads Mapped Confidently to Exonic Regions | 57.5% |
| Reads Mapped Confidently to Transcriptome | 54.3% |
| Reads Mapped Antisense to Gene | 1.1% |

##### Cells

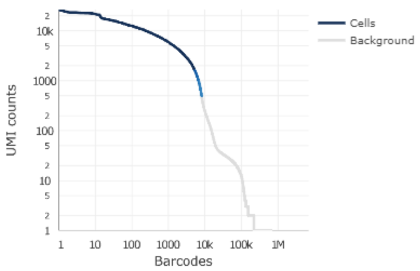


| | |
|---|---|
| Estimated Number of Cells | 6,937 |
| Fraction Reads in Cells | 88.0% |
| Mean Reads per Cell | 120,798 |
| Median Genes per Cell | 898 |
| Total Genes Detected | 14,493 |
| Median UMI Counts per Cell | 2,685 |

##### Sample

| | |
|---|---|
| Name | MLN2306 |
| Description | |
| Transcriptome | Bos_taurus.ARS-UCD1.2.dna.toplevel |
| Chemistry | Single Cell 3' v3 |
| Cell Ranger Version | 3.0.2 |



#### MLN3006

**The analysis detected some issues. Details »**

| Alert | Value | Detail |
|---|---|---|
| ⚠ **Low Fraction Valid UMIs** | 69.8% | Ideal > 75%. This may indicate a quality issue with the Illumina R2 read for Single Cell 3' v1 or the R1 read for Single Cell 3' v2/v3 and Single Cell 5'. Application performance may be affected. |

##### Estimated Number of Cells

5,187

| Mean Reads per Cell | Median Genes per Cell |
|---|---|
| 147,824 | 2,007 |

##### Sequencing

| | |
|---|---|
| Number of Reads | 766,767,159 |
| Valid Barcodes | 96.6% |
| Sequencing Saturation | 44.0% |
| Q30 Bases in Barcode | 96.3% |
| Q30 Bases in RNA Read | 93.9% |
| Q30 Bases in UMI | 96.2% |

##### Mapping

| | |
|---|---|
| Reads Mapped to Genome | 84.3% |
| Reads Mapped Confidently to Genome | 77.3% |
| Reads Mapped Confidently to Intergenic Regions | 16.9% |
| Reads Mapped Confidently to Intronic Regions | 26.2% |
| Reads Mapped Confidently to Exonic Regions | 34.2% |
| Reads Mapped Confidently to Transcriptome | 30.7% |
| Reads Mapped Antisense to Gene | 1.5% |

##### Cells

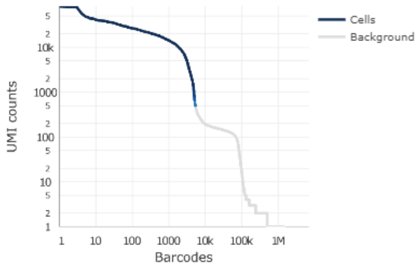


| | |
|---|---|
| Estimated Number of Cells | 5,187 |
| Fraction Reads in Cells | 79.1% |
| Mean Reads per Cell | 147,824 |
| Median Genes per Cell | 2,007 |
| Total Genes Detected | 15,844 |
| Median UMI Counts per Cell | 7,157 |

##### Sample

| | |
|---|---|
| Name | MLN3006 |
| Description | |
| Transcriptome | Bos_taurus.ARS-UCD1.2.dna.toplevel |
| Chemistry | Single Cell 3' v3 |
| Cell Ranger Version | 3.0.2 |

### Supplementary File 1A

SUMMARY ANALYSIS

#### MLN0707

The analysis detected some issues. Details »

| Alert | Value | Detail |
|---|---|---|
| Low Fraction Valid UMIs | 69.8% | Ideal > 75%. This may indicate a quality issue with the Illumina R2 read for Single Cell 3' v1 or the R1 read for Single Cell 3' v2/v3 and Single Cell 5'. Application performance may be affected. |

##### Estimated Number of Cells

4,288

| Mean Reads per Cell | Median Genes per Cell |
|---|---|
| 176,752 | 2,155 |

##### Sequencing

| | |
|---|---|
| Number of Reads | 757,914,774 |
| Valid Barcodes | 96.5% |
| Sequencing Saturation | 49.9% |
| Q30 Bases in Barcode | 96.3% |
| Q30 Bases in RNA Read | 94.1% |
| Q30 Bases in UMI | 96.2% |

##### Mapping

| | |
|---|---|
| Reads Mapped to Genome | 85.8% |
| Reads Mapped Confidently to Genome | 79.6% |
| Reads Mapped Confidently to Intergenic Regions | 16.7% |
| Reads Mapped Confidently to Intronic Regions | 27.5% |
| Reads Mapped Confidently to Exonic Regions | 35.4% |
| Reads Mapped Confidently to Transcriptome | 32.0% |
| Reads Mapped Antisense to Gene | 1.5% |

##### Cells

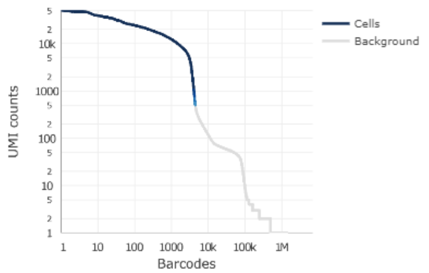


| | |
|---|---|
| Estimated Number of Cells | 4,288 |
| Fraction Reads in Cells | 86.9% |
| Mean Reads per Cell | 176,752 |
| Median Genes per Cell | 2,155 |
| Total Genes Detected | 15,713 |
| Median UMI Counts per Cell | 7,819 |

##### Sample

| | |
|---|---|
| Name | MLN0707 |
| Description | |
| Transcriptome | Bos_taurus.ARS-UCD1.2.dna.toplevel |
| Chemistry | Single Cell 3' v3 |
| Cell Ranger Version | 3.0.2 |

**Supplementary File 1A** Cell Ranger summaries for the three samples (MLN2306, MLN3006, MLN0707) analyzed by 10x scRNA-seq.

### Supplementary File 1B

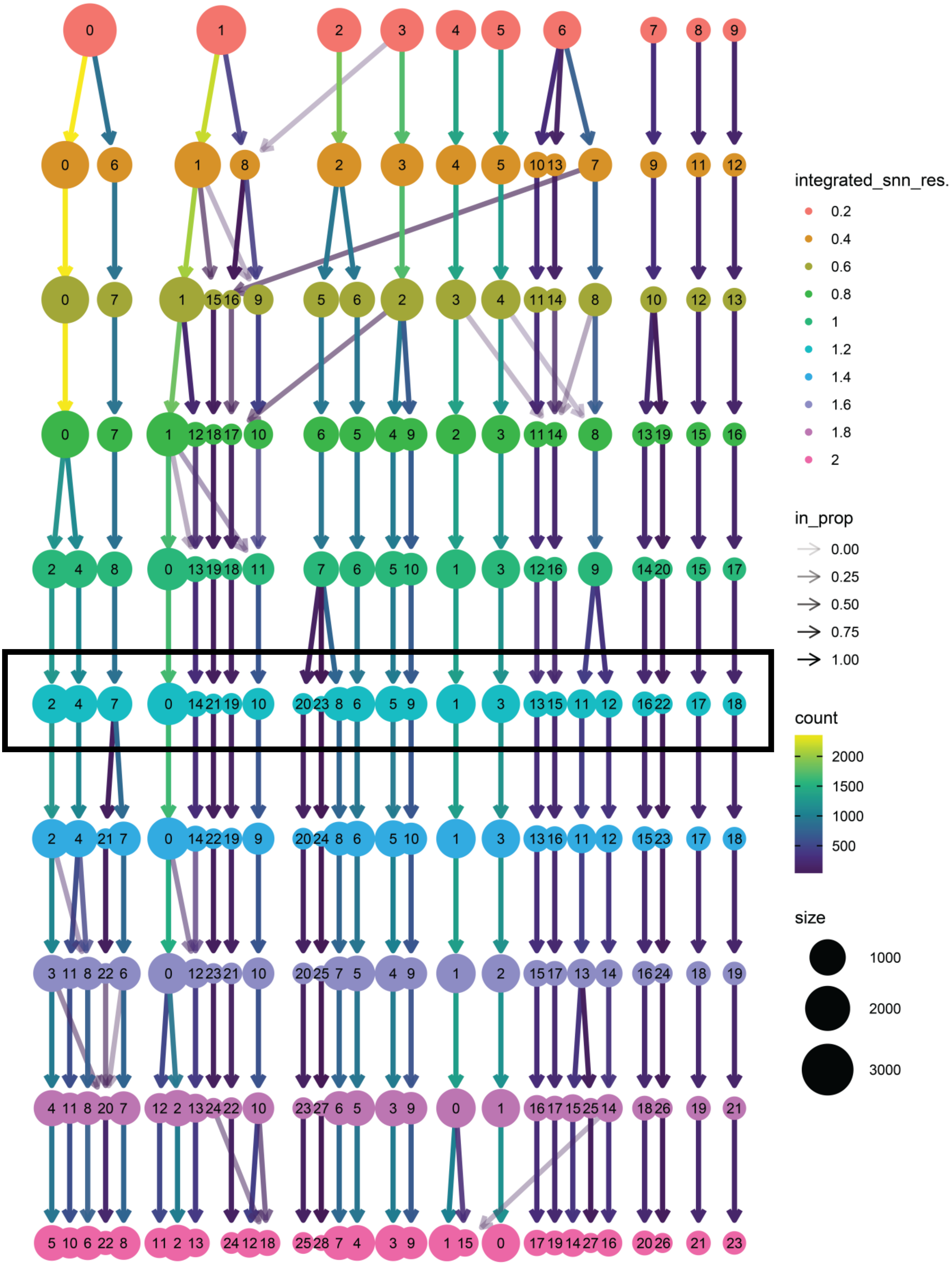


**Supplementary File 1B** Evaluation of cluster stability (integrated dataset) using the clustree package (Zappia et al. 2018). A resolution of 1.2 was chosen for downstream analyses, resulting in 24 different clusters.
