## Supplementary material for "Single-cell transcriptomics reveals striking heterogeneity and functional organization of dendritic and monocytic cells in the bovine mesenteric lymph node": Suppl_5_keyGenes

### Supplementary File 5

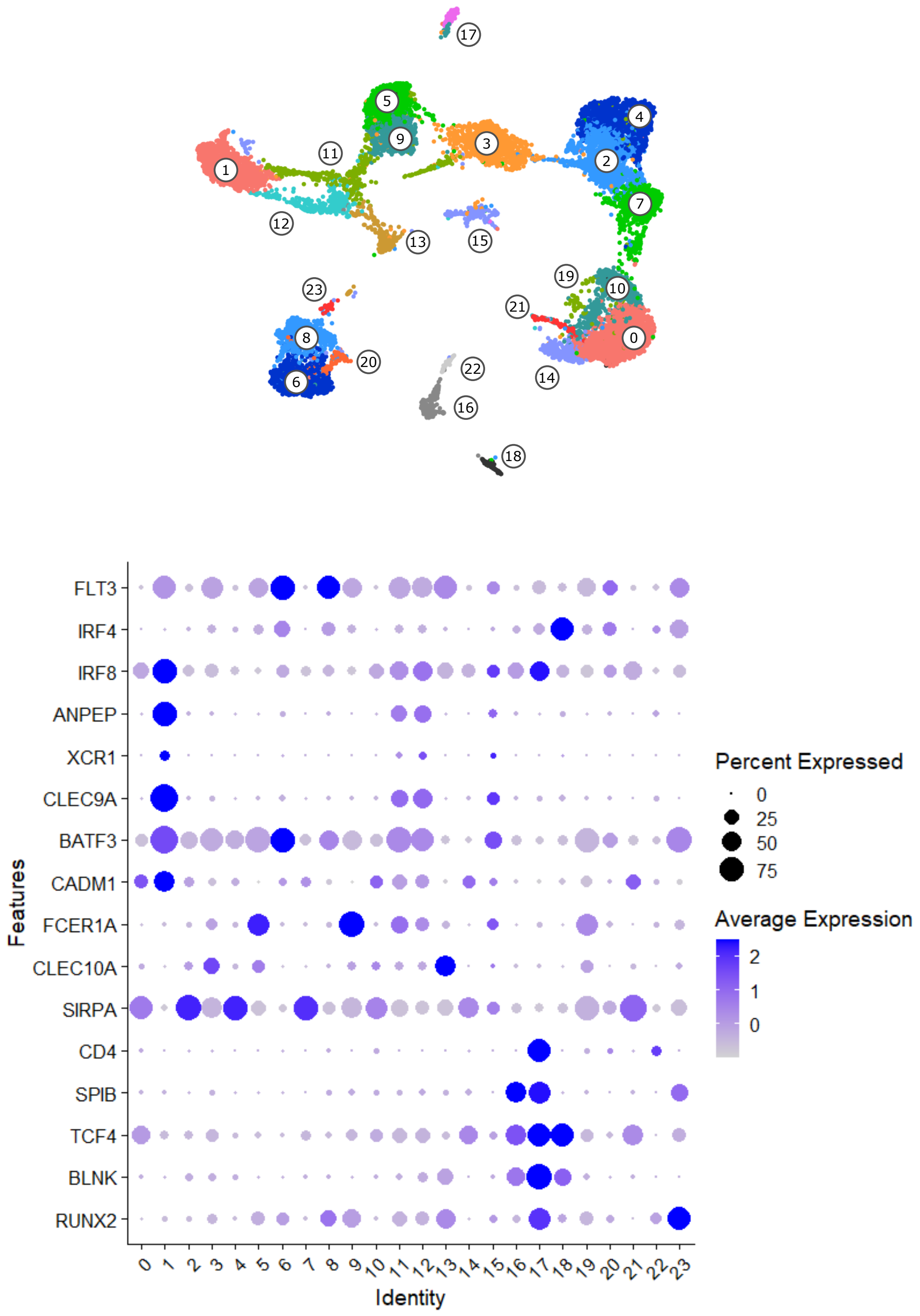


**Supplementary File 5** Key-gene expression defining DC subsets visualized in a dot plot for the complete dataset.
