## Supplementary material for "Single-cell transcriptomics reveals striking heterogeneity and functional organization of dendritic and monocytic cells in the bovine mesenteric lymph node": Suppl_12_GOI

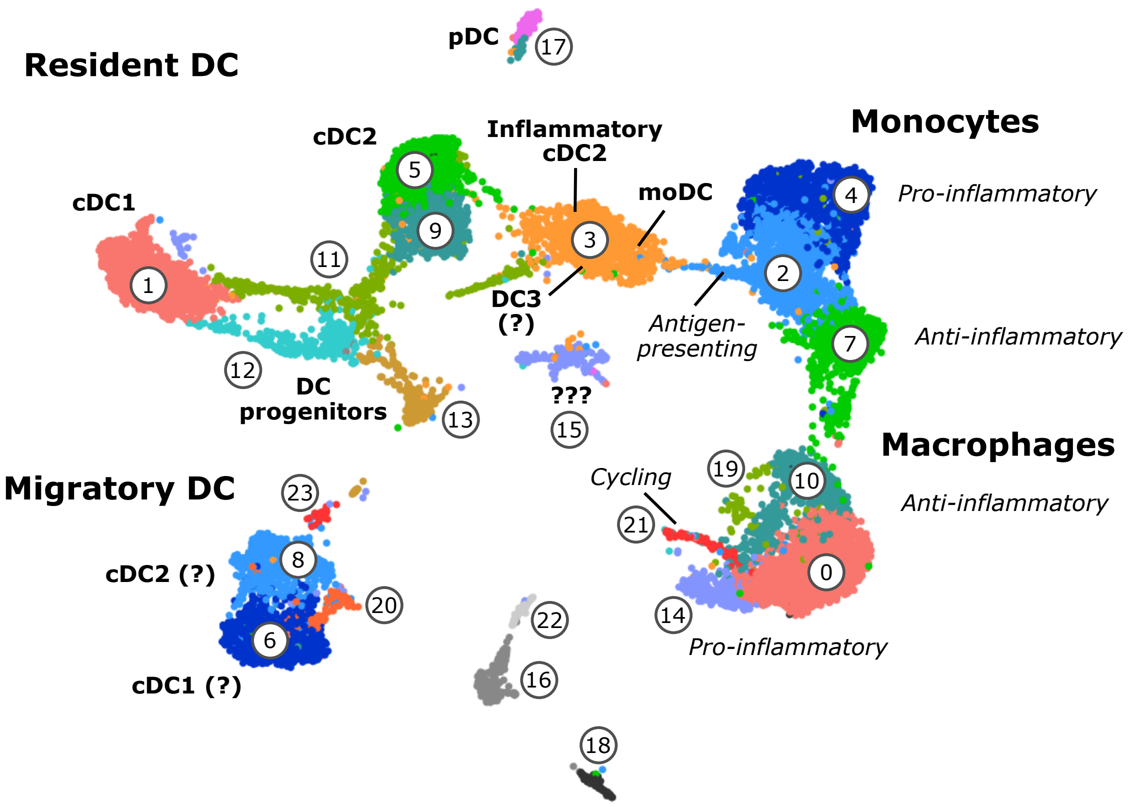


### Visualization of selected genes

**Page 2**
1) Pattern-recognition receptors
2) Fc receptors
3) Purinergic receptors

**Page 3**
4) Chemokines
5) Chemokine receptors

**Page 4**
6) Integrins
7) Galectins

**Page 5**
8) Antigen presentation
9) T-cell modulation

**Page 6**
10) Interleukins
11) Interleukin receptors

**Page 7**
12) TNF
13) TNF receptors

**Page 8**
14) Tetraspanins
15) Metalloproteinases

**Page 9**
16) Metabolism (misc.)
17) Glycolysis

**Page 10+11**
18) Solute carriers

**Page 12**
19) Complement system
20) Interferon-associated

**Page 13**
21) Retinoic-acid production and signaling
22) Semaphorins and receptors

#### 1) Pattern-recognition receptors

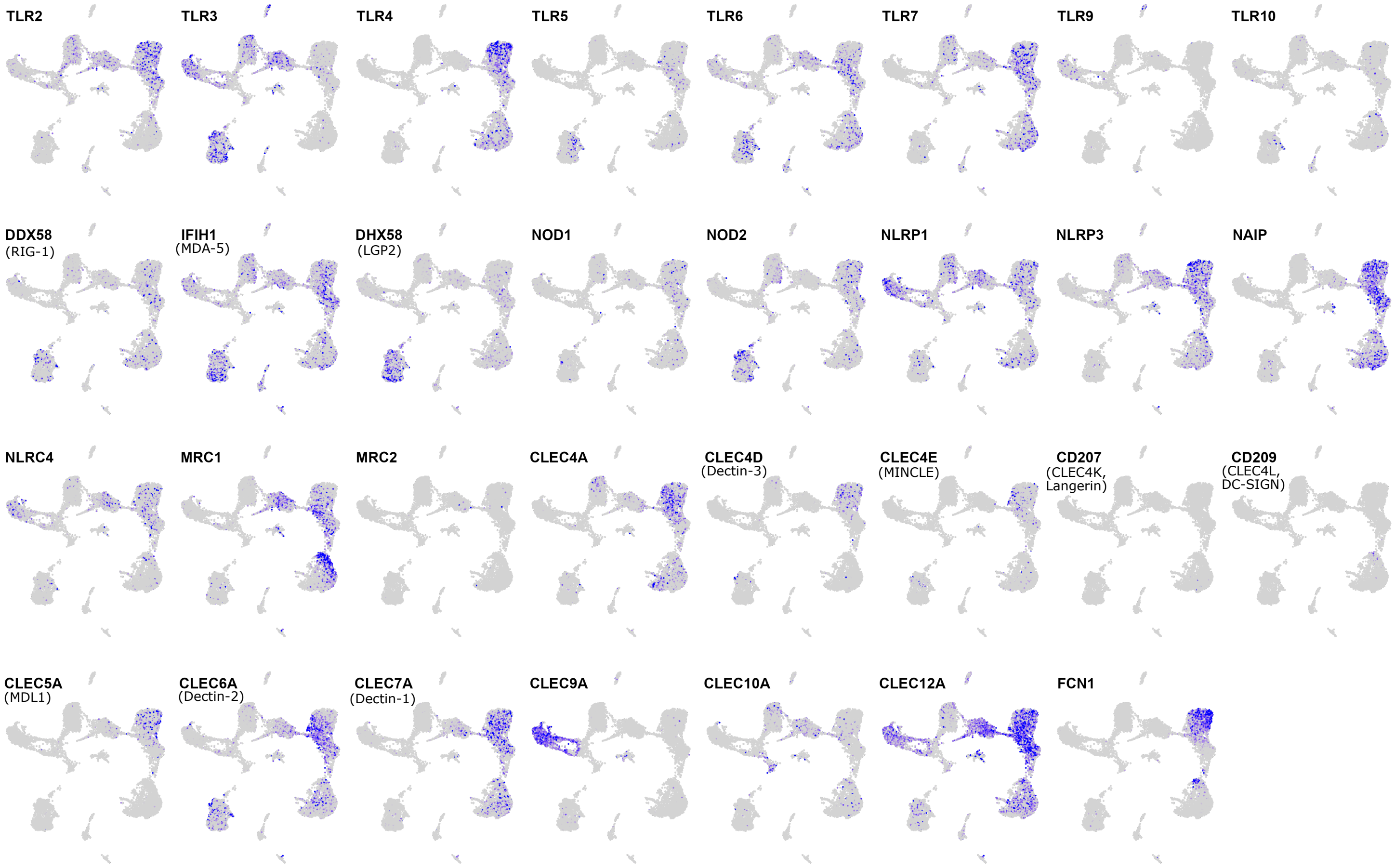


#### 2) Fc receptors

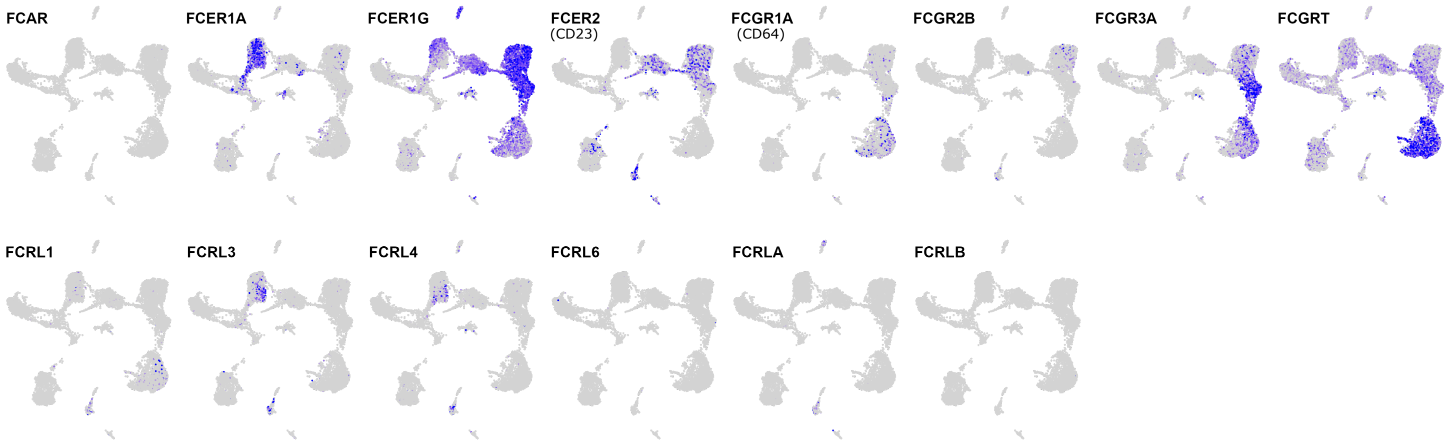


#### 3) Purinergic receptors

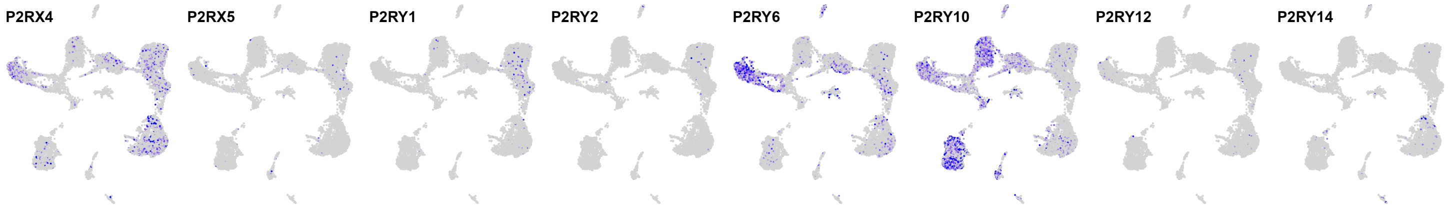


Poorly detected: P2RX1, P2RX2, P2RX3, P2RX7, P2RY8, P2RY11

#### 4) Chemokines

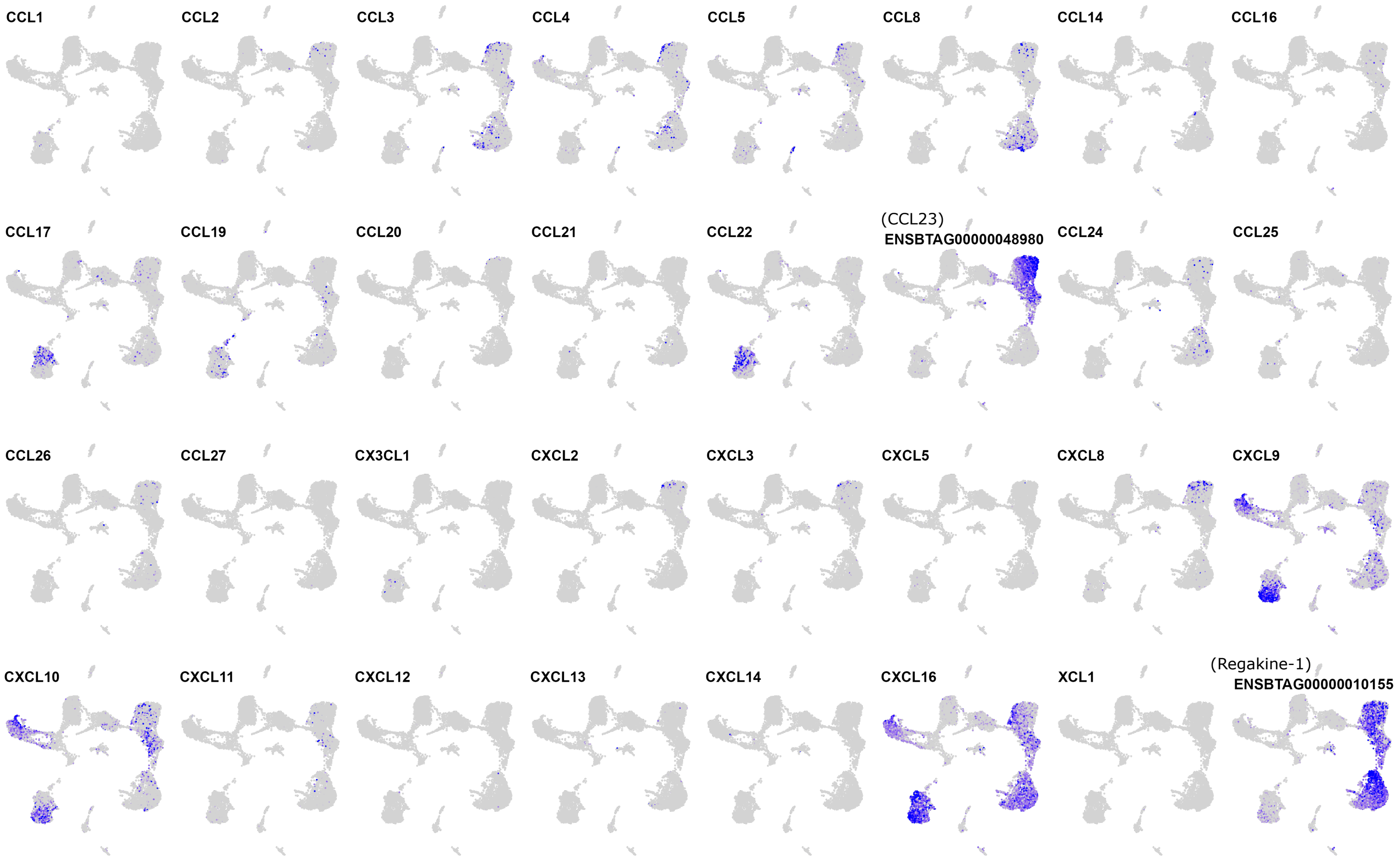


#### 5) Chemokine receptors

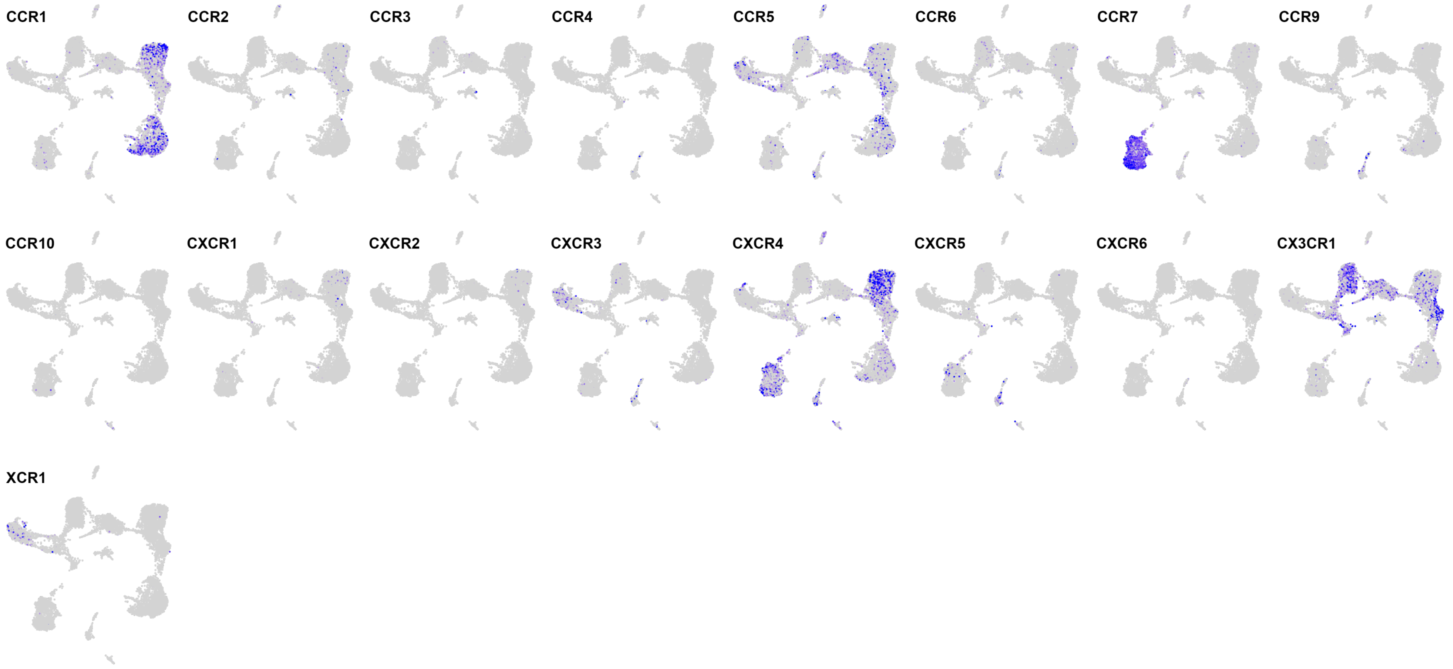

#### 6) Integrins

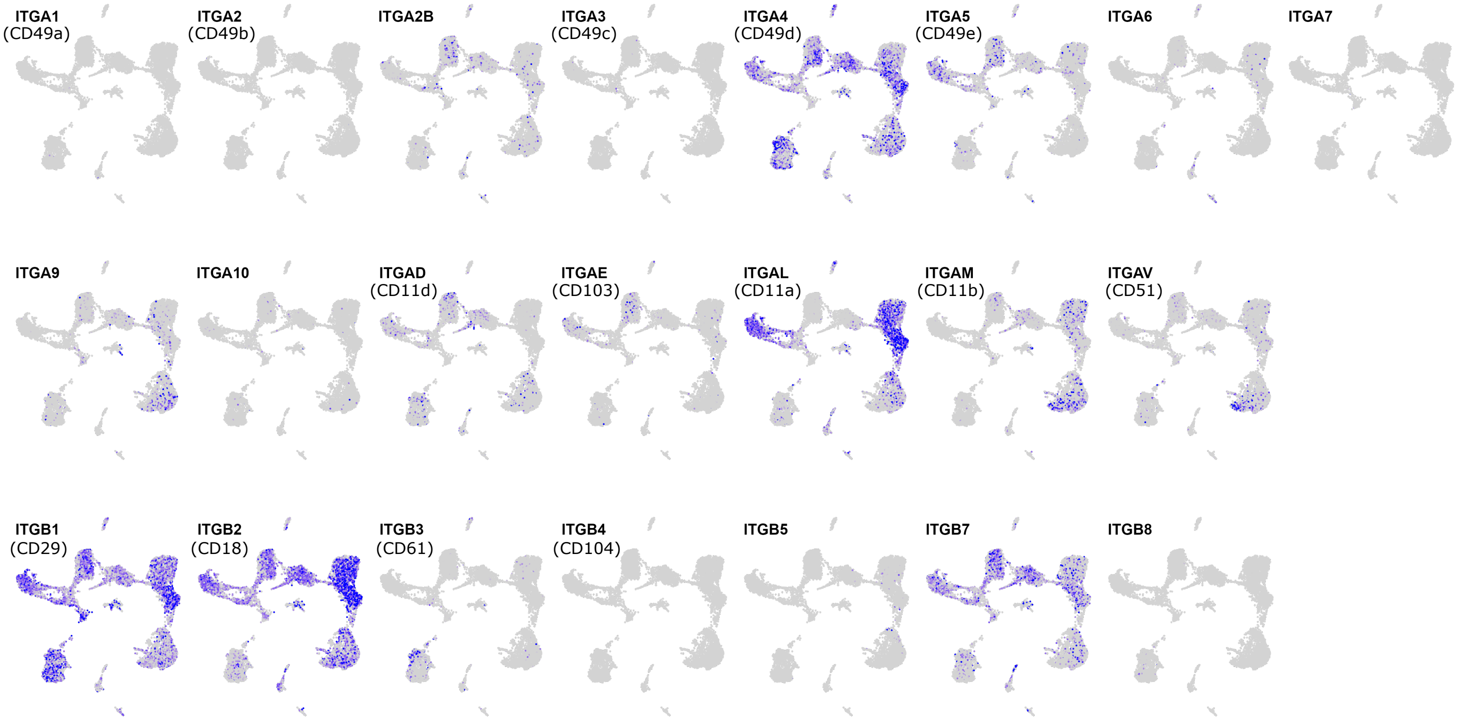


#### 7) Galectins

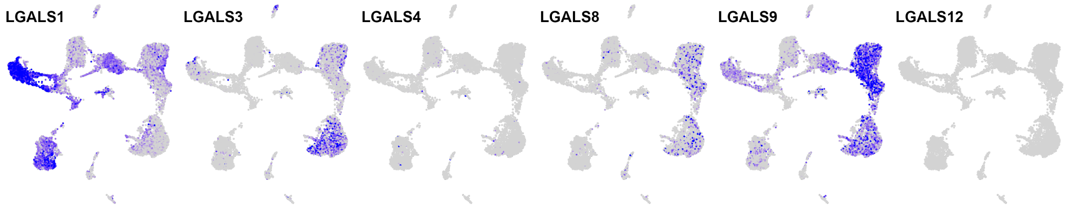

#### 8) Antigen presentation

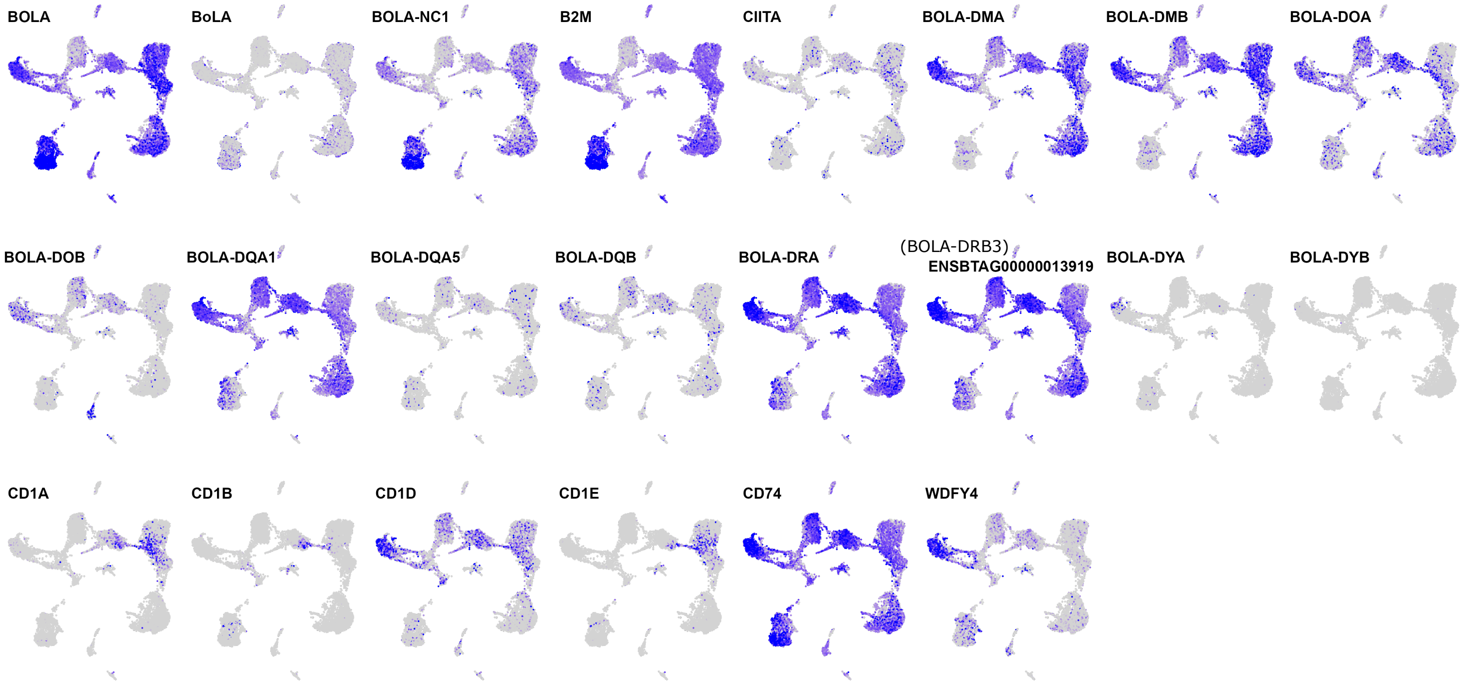


#### 9) T-cell modulation

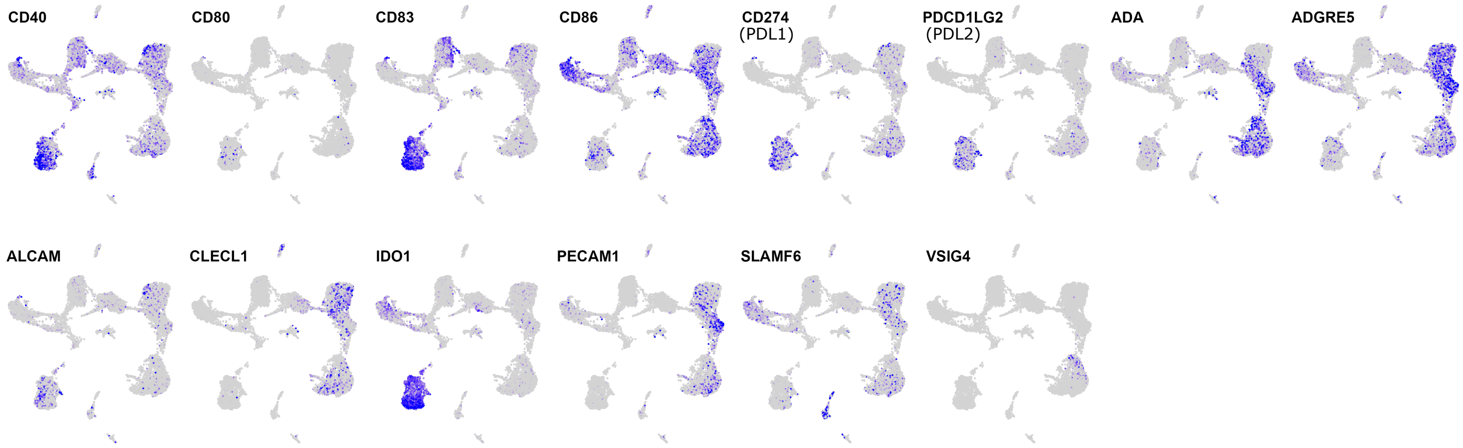

#### 10) Interleukins

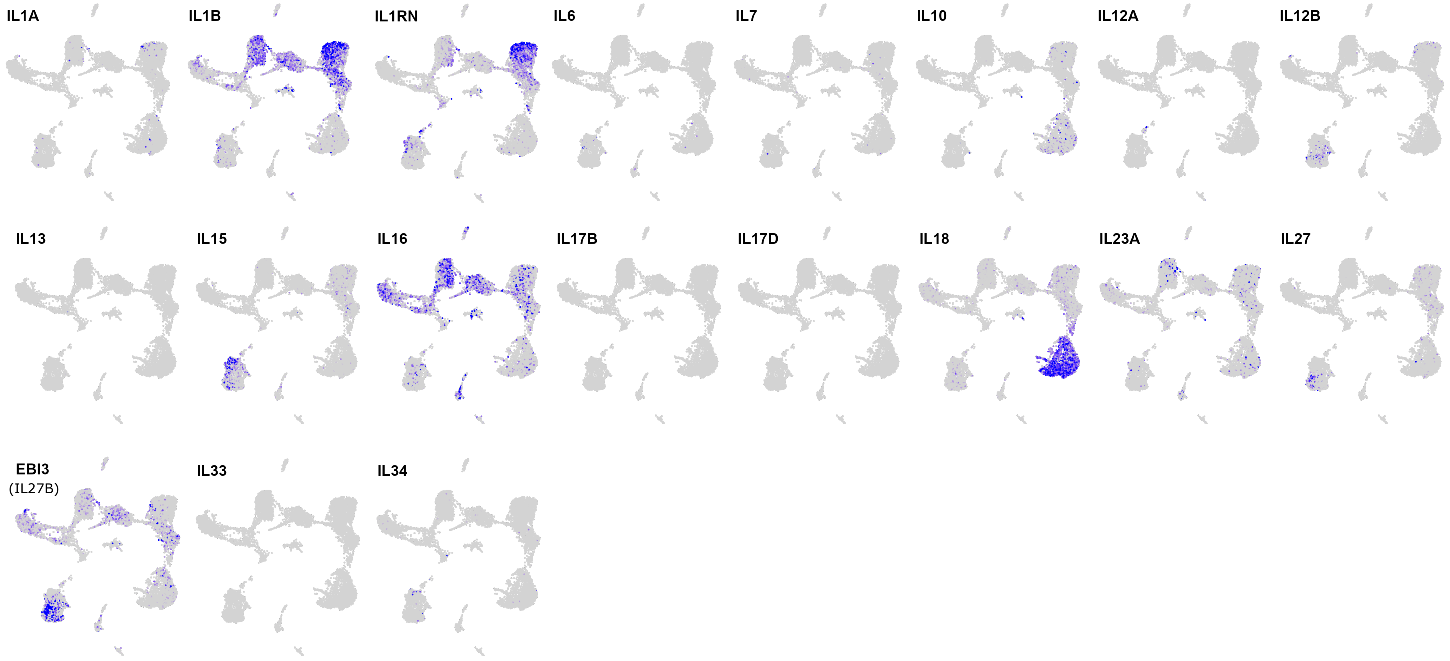


#### 11) Interleukin receptors

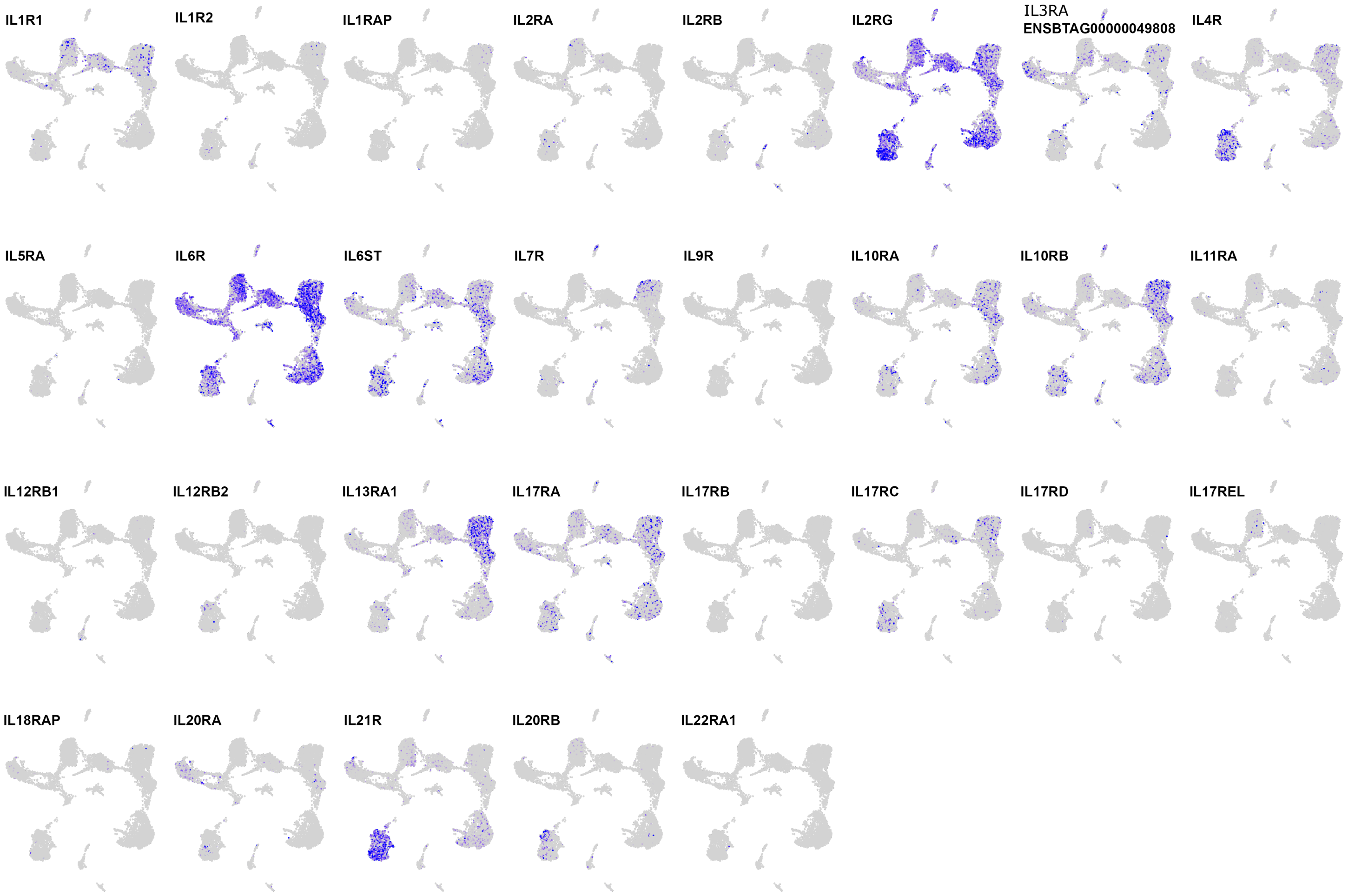

#### 12) TNF superfamily

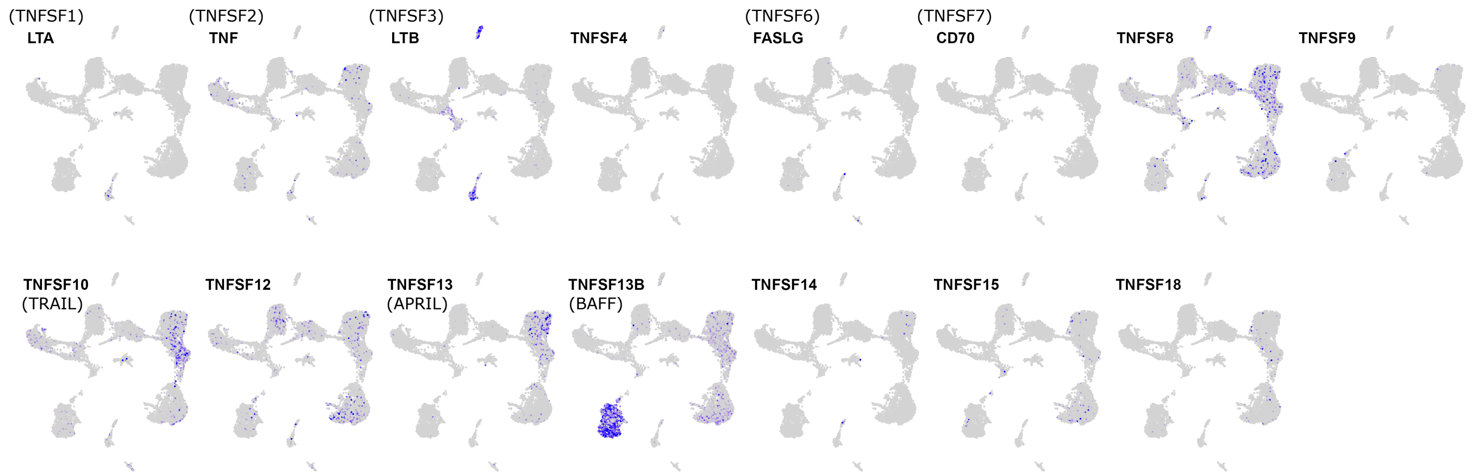


#### 13) TNF receptor superfamily

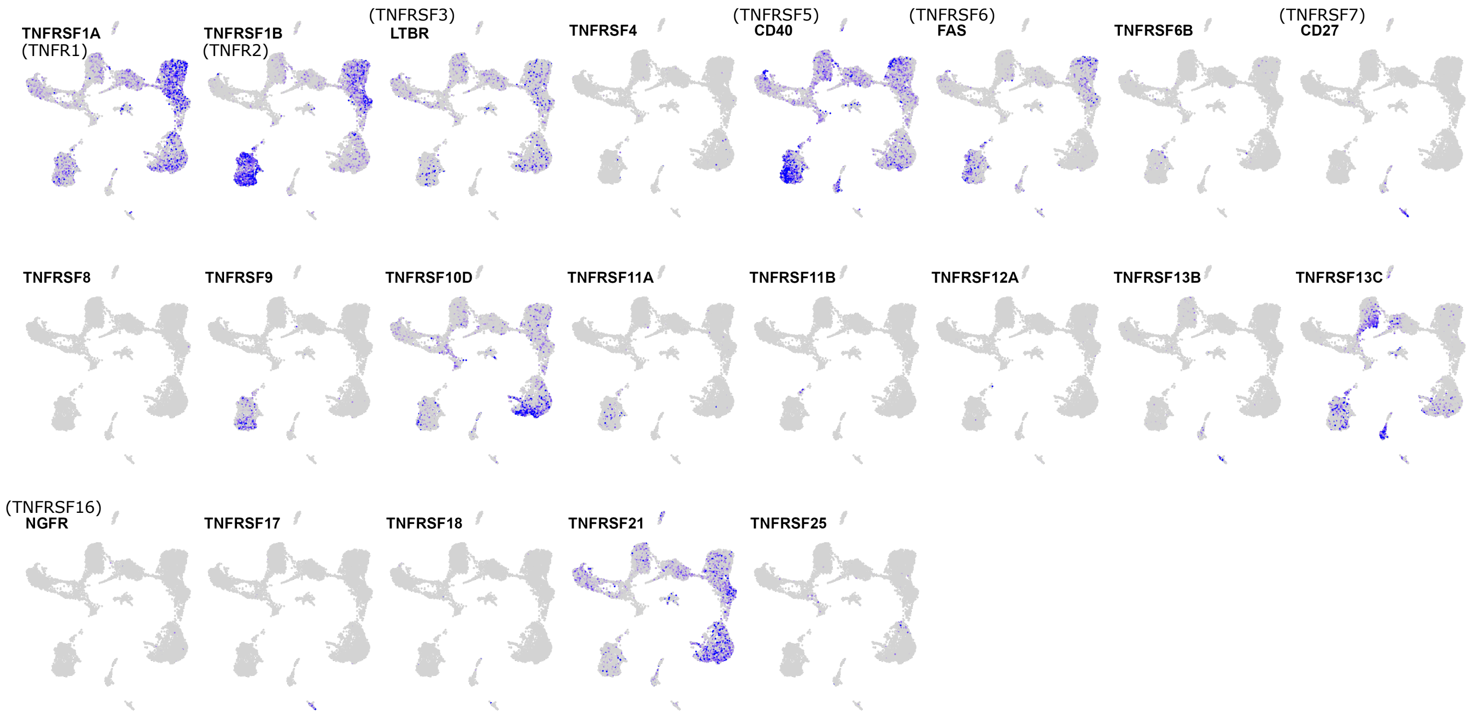

#### 14) Tetraspanins

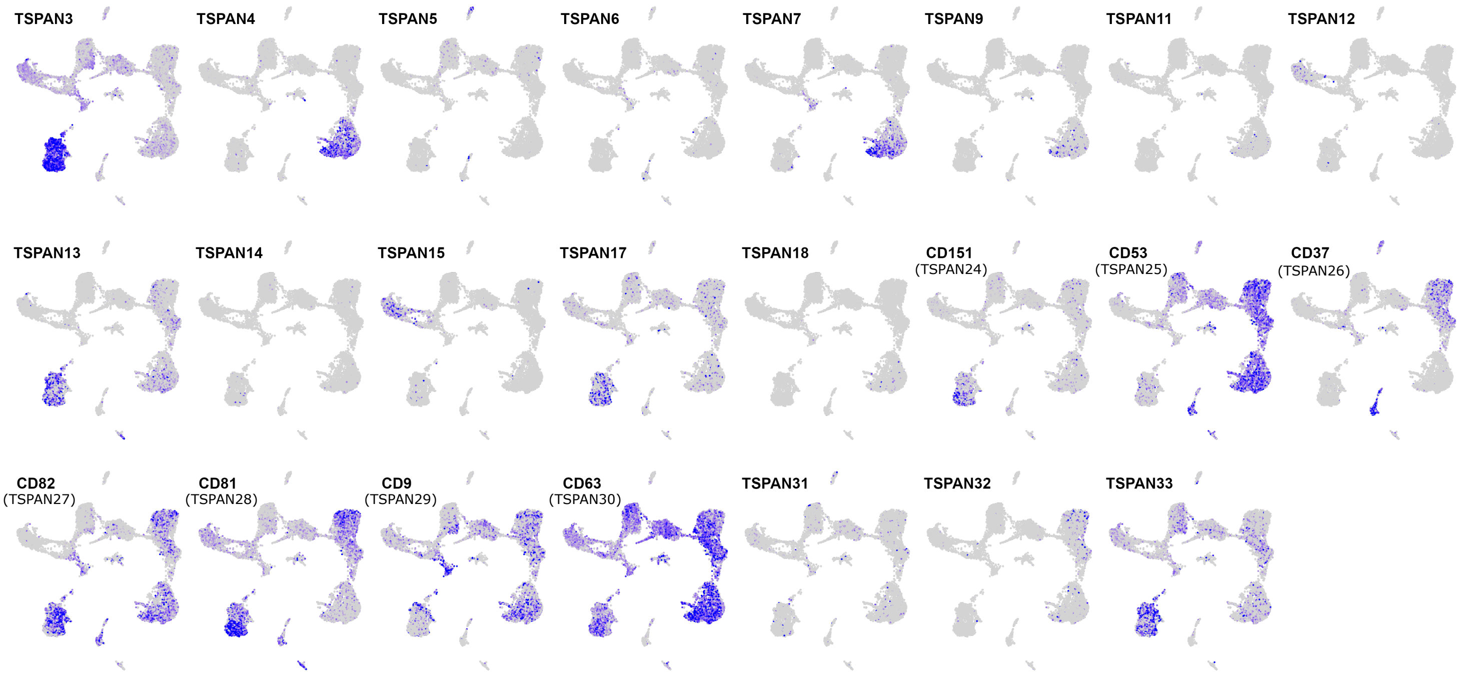


Poorly detected: TSPAN1, TSPAN2, UPK1B (TSPAN20)

#### 15) Metalloproteinases

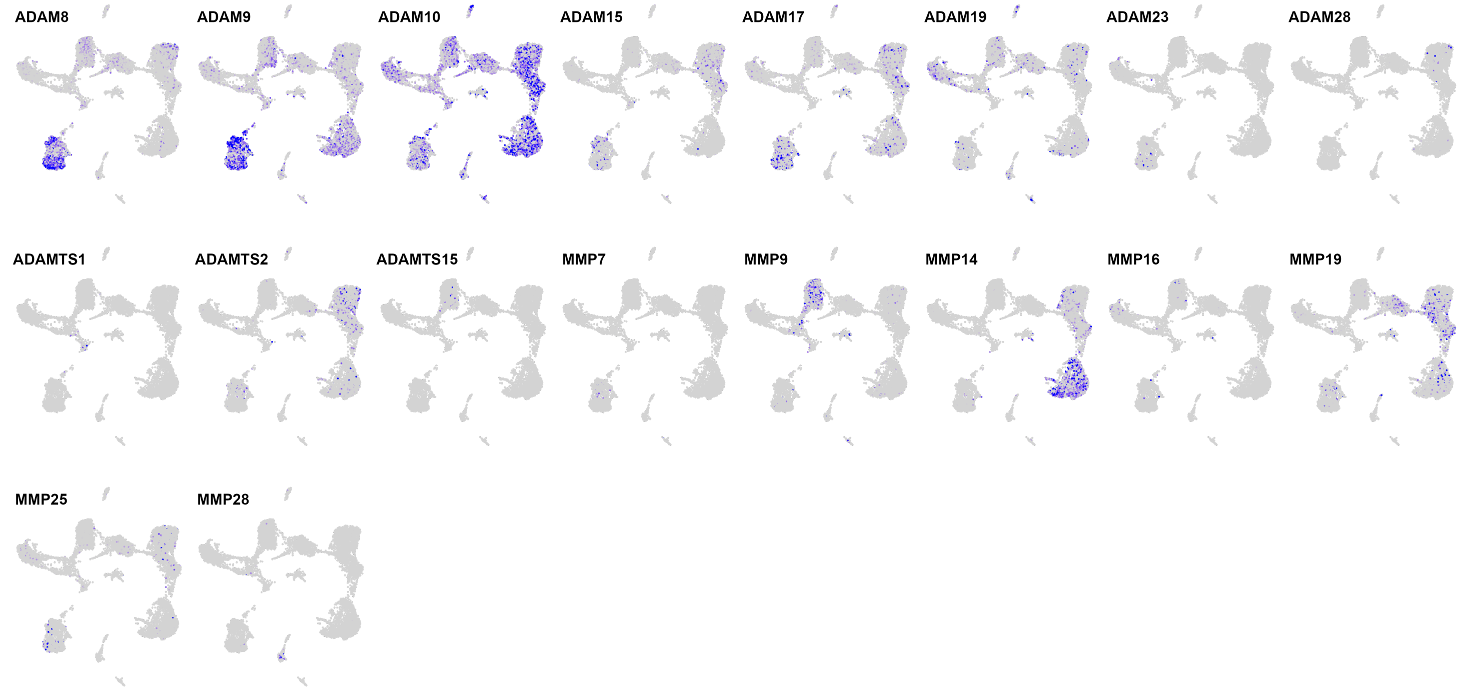


Poorly detected: ADAM11, ADAM12, ADAM20, ADAM22, ADAM32, ADAM33, ADAMTS6, ADAMTS7, ADAMTS8, ADAMTS10, ADAMTS12, ADAMTS14, MMP12, MMP15, MMP20

#### 16) Metabolism (misc.)

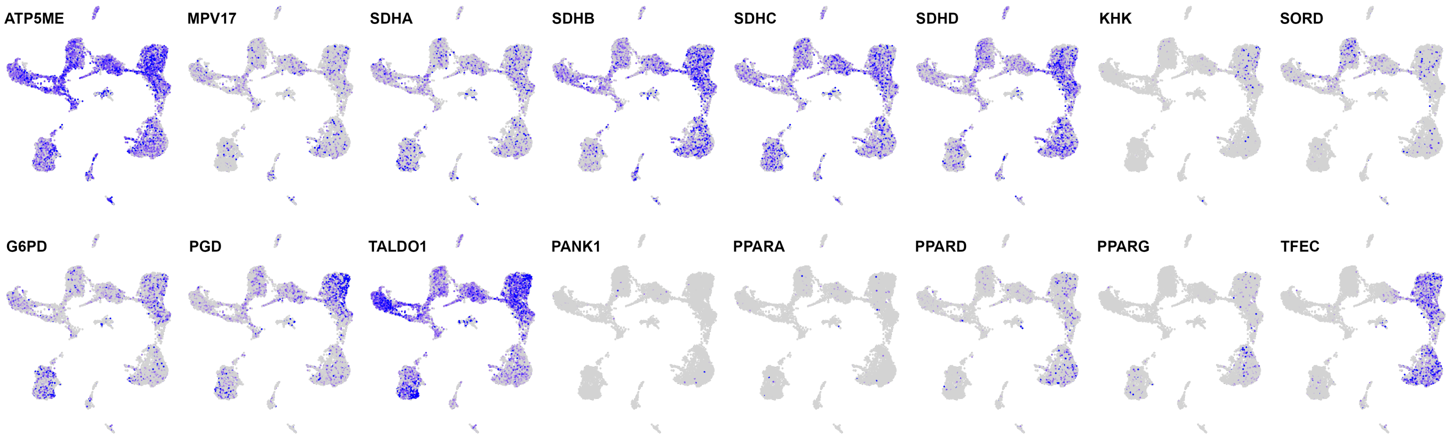


#### 17) Glycolysis

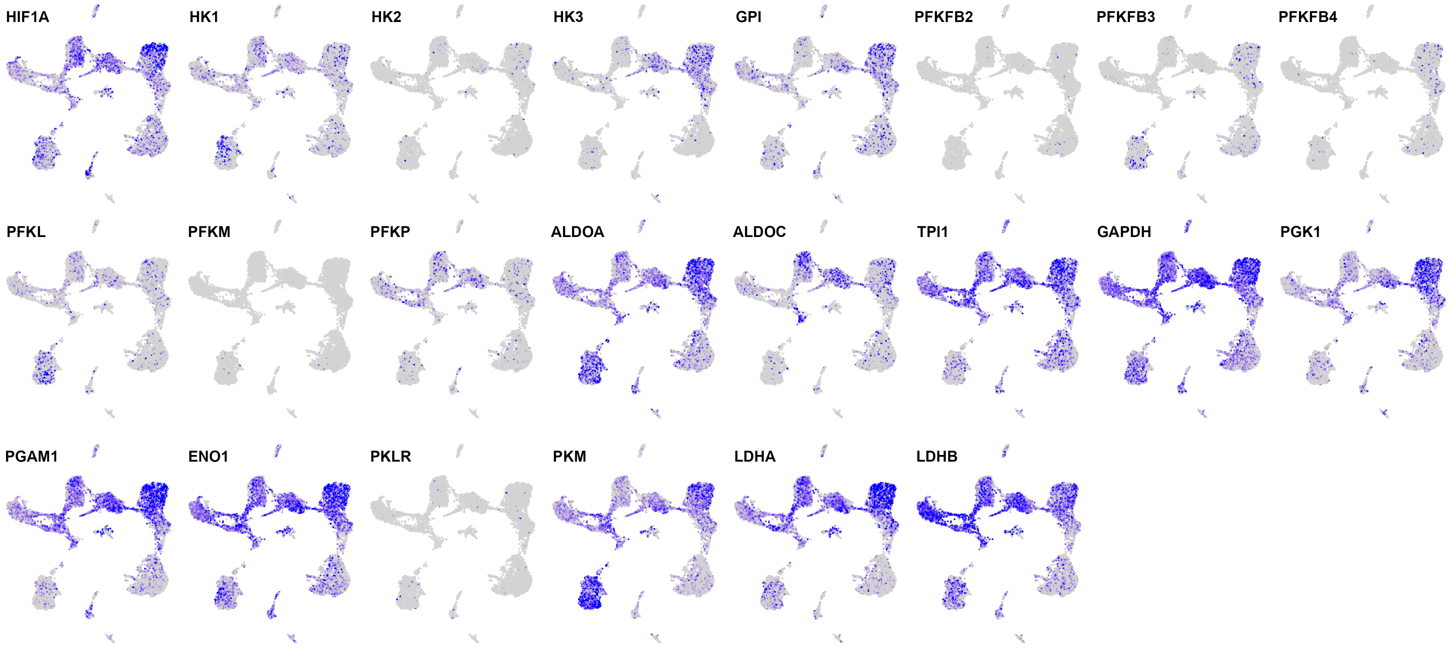

#### 18) Solute carriers (part 1/2)

##### Sugars

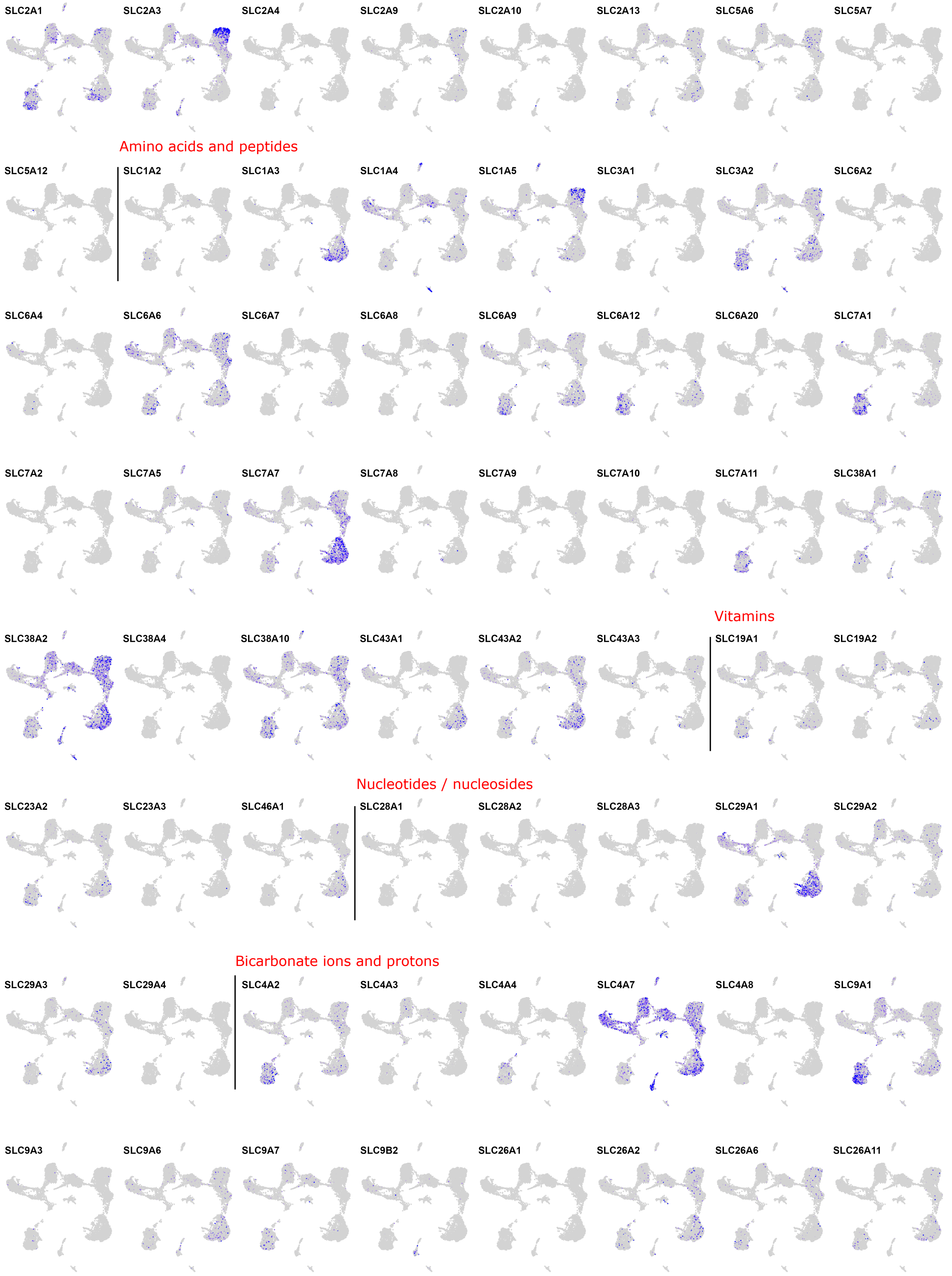

### 18) Solute carriers (part 2/2)

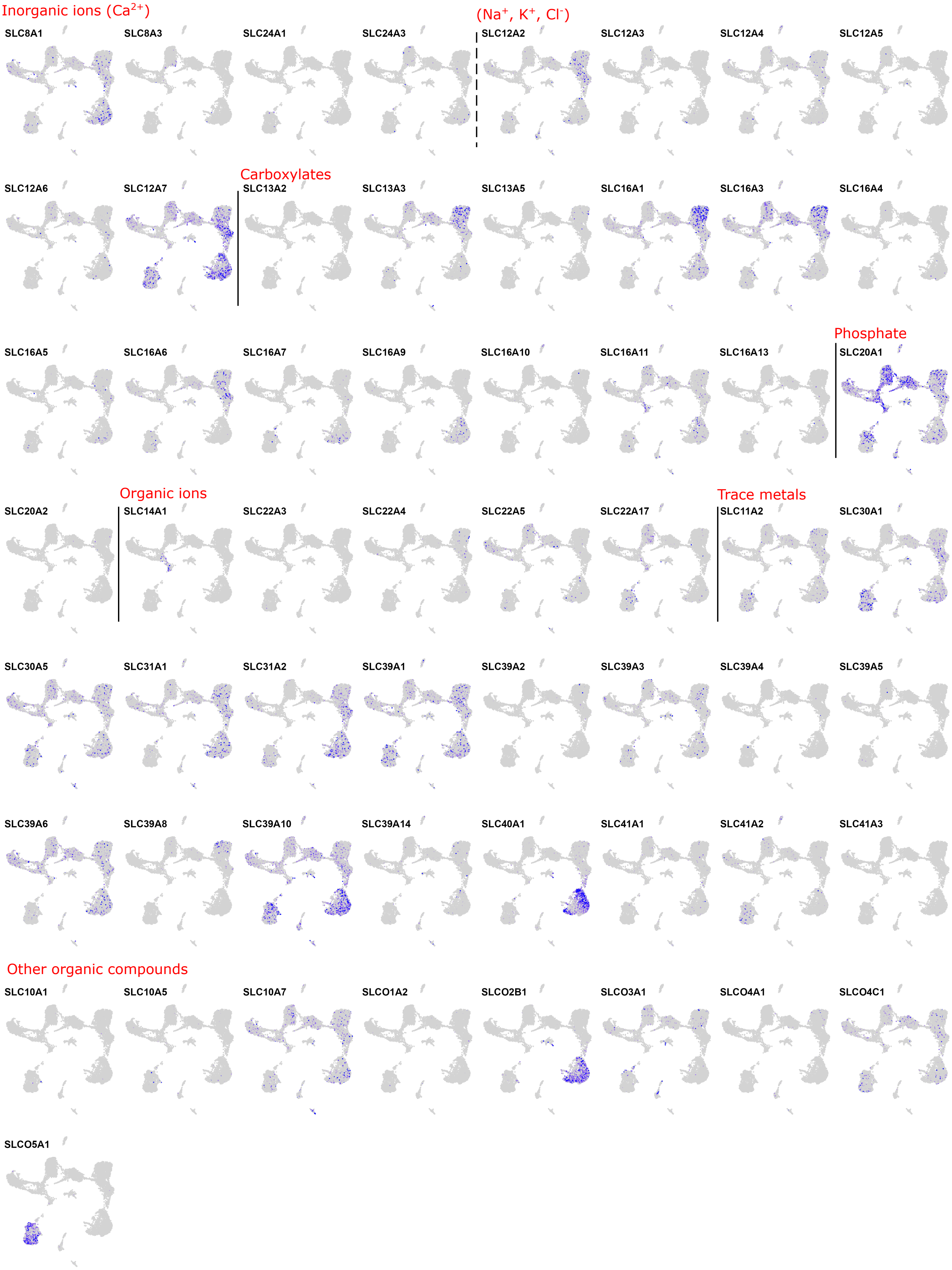

#### 19) Complement system

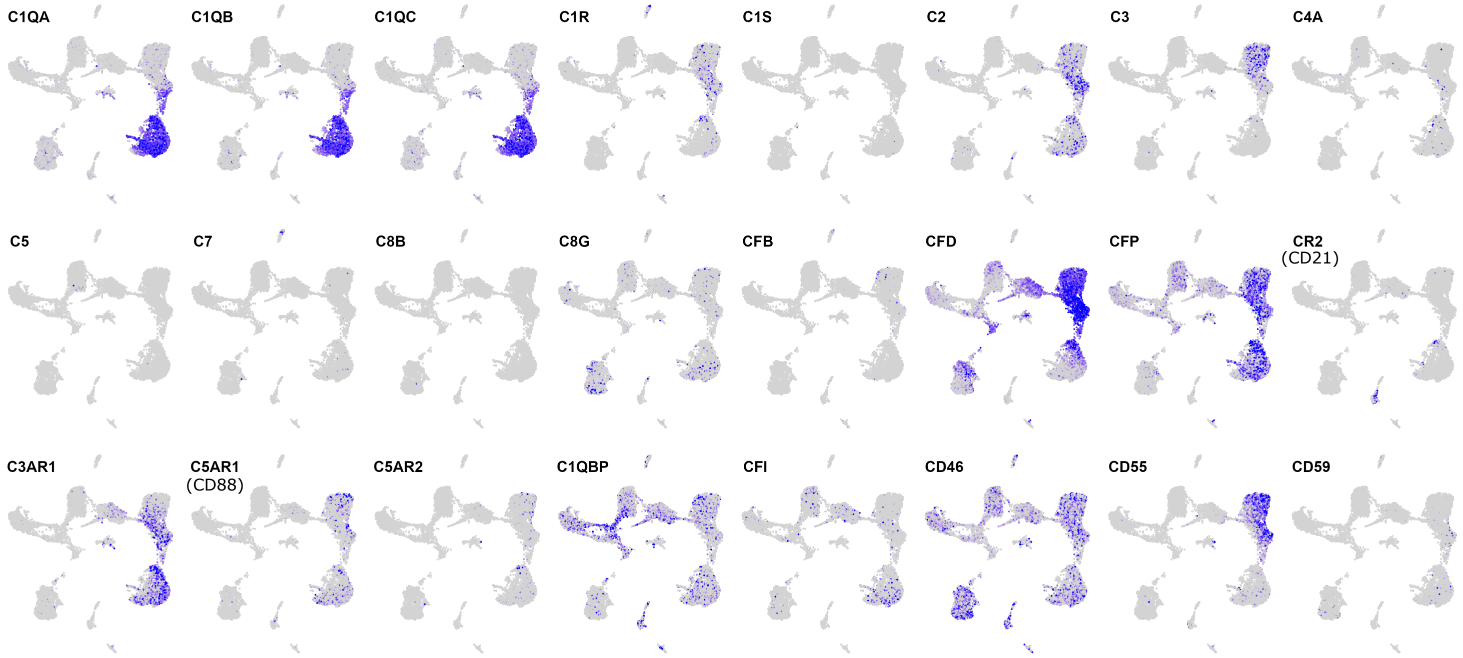


#### 20) Interferon-associated

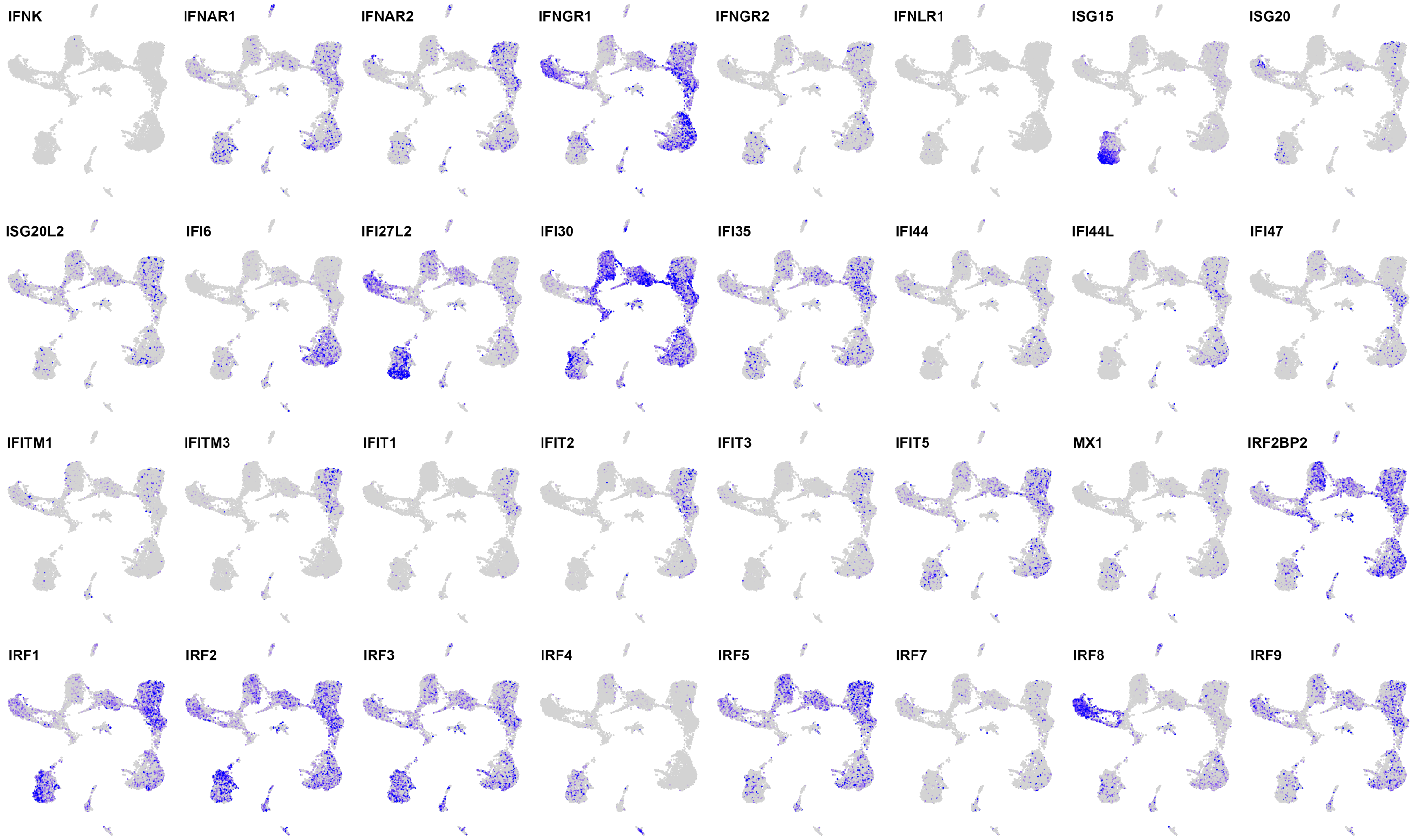

#### 21) Retinoic-acid production and signaling

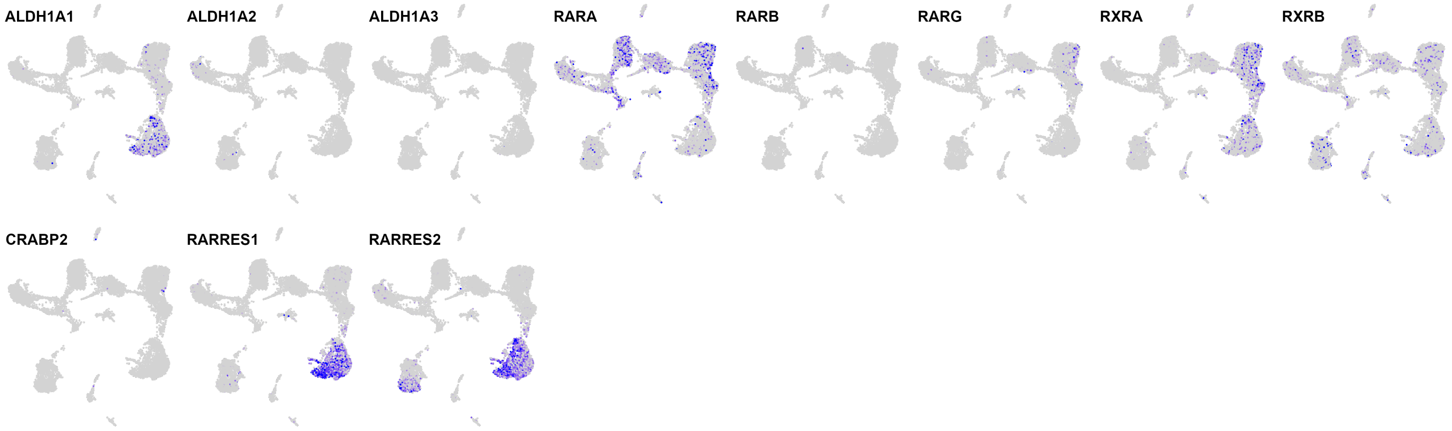


#### 22) Semaphorins and receptors

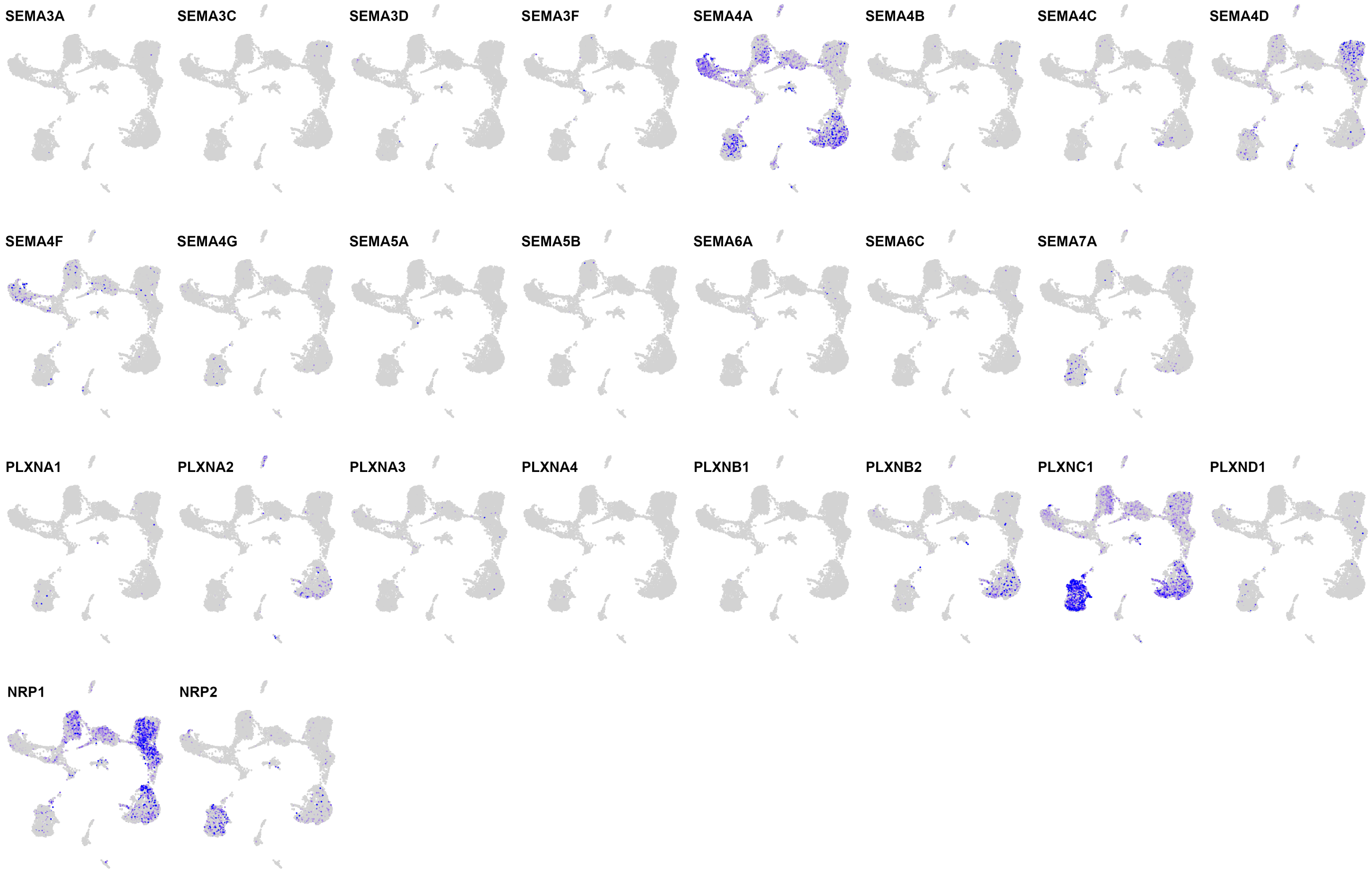
